# Microbial communities demonstrate robustness in stressful environments due to predictable composition shifts

**DOI:** 10.1101/2025.08.21.667681

**Authors:** Jana S. Huisman, Martina Dal Bello, Jeff Gore

**Author notes:** Correspondence (J.S. Huisman).

## Abstract

Environmental stress reduces species growth rates, but its impact on the function of microbial communities is less clear. Here, we experimentally demonstrate that increasing salinity stress shifts community composition towards species with higher growth rates. As a result, the mean community growth rate is more robust to increasing stress than the growth of individual species. We show this by propagating natural aquatic communities at multiple salinities and mapping the observed diversity onto the measured salinity performance curves of *>*80 bacterial isolates. We further validate these results with pairwise species competitions and in metagenomic data of natural communities sampled from estuarine environments. A Lotka-Volterra model including mortality and salinity-dependent growth rates recapitulates the observed robustness of community growth sustained by more abundant faster growers at high salinity. These results extend to other environmental stressors and point to fundamental mechanisms with which communities maintain growth despite deteriorating conditions.

## Introduction

Microbial communities fulfill crucial ecosystem functions across soil, water and host-associated environments, from nutrient cycling to pathogen suppression [1, 2]. These functions are emergent, community-level properties resulting from the physiology of individual community members, their relative abundances, and the interactions between them [3]. Simple perturbations that affect individual species can have complex, nonlinear effects on the community. As a result, it remains unclear whether community-level properties are predictable in changing environments. Specifically, we ask how environmental stress – an abiotic change which reduces the growth of most species – impacts community composition and function.

Recent work has demonstrated that environmental changes can have predictable effects on community composition, although the functional consequences are less clear. Increasing temperature – which increases growth rates of most species – selects for communities dominated by slow growing species, both in controlled laboratory experiments with two or three species [4] and in marine microbial communities assayed with metagenomic data [5]. At the same time, increased population-wide mortality (imposed through dilution) favors the faster growing species in pairwise competitions [6]. However, the impact of such compositional shifts on community function – and how this depends on the specific environmental change – remains understudied [7]. An environmental stressor may impair community function as it reduces species growth rates, and the resulting shifts in the community composition could both amplify or buffer this effect.

The most fundamental functions one can ascribe to a microbial community are the ability to grow and produce biomass. This forms the basis for all other roles microbes play in an ecosystem. For instance, the rate of biomass production (the ‘productivity’) has been shown to predict invasion resistance against new species introductions [8]. The mean community growth rate becomes especially important in perturbed environments as it describes the rate at which the community recovers from the perturbation. In addition, communities may perform specific functions that are relevant to humans, animals, or ecosystems (e.g. producing the characteristic taste of chocolate [9], or degrading organic carbon and producing methane [10, 11]). In the following, we will call a community-level functional trait ‘robust’ if this trait is less affected by perturbation than expected based on the mean behavior of the individual species.

Salinity is a key environmental stressor structuring microbial communities, particularly in soil and aquatic environments [12]. Climate change is expected to cause major disruption to existing environmental salinity gradients. Sea-level rise, anthropogenic land use changes, and shifting precipitation patterns are increasing both the extent and variability of salt intrusion into freshwater systems [13, 14]. These salinity shifts can restructure microbial communities in complex ways: in salt marshes, for example, salinization reduces microbial methanogenesis and increases sulfur reduction [15]. Observations abound of community composition changing with salinity [12, 16, 17, 18, 19, 20, 21], yet we lack the ability to predict the strength and direction of this compositional shift and its functional consequences.

In contrast to the community-level ecological response, the mechanistic effects of salinity on individual bacteria are quite well understood. Salinity increases the osmolarity of the medium, causing water to flow out of bacterial cells and their turgor pressure to drop [22]. This physiological stress reduces growth of most microbes [22]. The magnitude of this growth reduction is linked to species-specific mechanisms of osmoadaptation, which also determine the salinity at which a species achieves optimal growth [22, 23]. While this optimal salinity is higher for salt-adapted (halotolerant or halophilic) bacteria than for other species, the mechanisms of stress beyond the optimal salinity are likely similar. However, a quantitative exploration of the relation between growth and salinity has remained restricted to common foodborne pathogens [24, 25], and little is known for environmental microbes. Moreover, the effects of salinity on community-level properties such as carrying capacity, competitive interactions, and overall growth remain largely unquantified.

Here, we ask to which extent community-level responses to increasing salinity can be predicted from traits of the constituent species. We propagated natural communities at multiple salinities, and found that the mean community growth rate was remarkably robust to an increase in salinity despite decreasing growth rates for individual species. Combining measured salinity performance curves – the maximum growth rate of a species as a function of salinity – with a general model of bacterial competition under environmental stress, we could explain that the robustness of the community growth rate is the result of a shift in community composition towards species with high growth rates in stressful environments. We validated these predictions with pairwise competitions and metagenomic data of communities sampled across environmental salinity gradients. Since we predict that any environmental stressor that reduces the growth rate of most species will lead to these composition shifts and the associated robustness, this points towards fundamental mechanisms by which communities maintain function in stressful environments.

## Results

### Community growth rates are robust due to compositional changes caused by environmental stress

To study how natural communities respond to increased environmental stress, we monitored the assembly of aquatic microbial communities at three different salinities (16, 31, 46 g/L sea salts), with fixed nutrient concentration and proportions of the five primary sea salts (NaCL, MgCl_2_, MgSO_4_, CaCl_2_, KCl). We sampled four aquatic microbial communities along a salinity gradient around the Boston harbor: the Charles River at the MIT sailing pavilion (“Brackish”; 4 g/L), the Boston harbor near the Institute for Contemporary Art (“Estuary”; 30 g/L), and the ocean at Canoe Beach, Nahant (“Marine 1 & 2”; 35 g/L; Fig. 1A). We serially propagated these communities *in vitro* every 2 days for a total of 7 cycles and assessed the diversity and composition of communities at 6 time points via 16S amplicon sequencing. As expected, some species could not grow under these laboratory conditions and the diversity of the environmental communities decreased in the first 1-2 cycles, across all propagation salinities (Fig. S1). By the 7th cycle, the majority of the communities reached a stable composition with 50-100 unique ASVs (Fig. S2, S3, S4). Across the starting communities and propagation salinities, notably different steady state compositions were reached (Fig. S5). While the species richness and Shannon diversity decreased at higher salinity (Fig. 1B, S1), community biomass remained remarkably constant (Fig. S6). This raises the question whether we can predict changes in community composition in response to increasing salinity.

**Figure 1:**
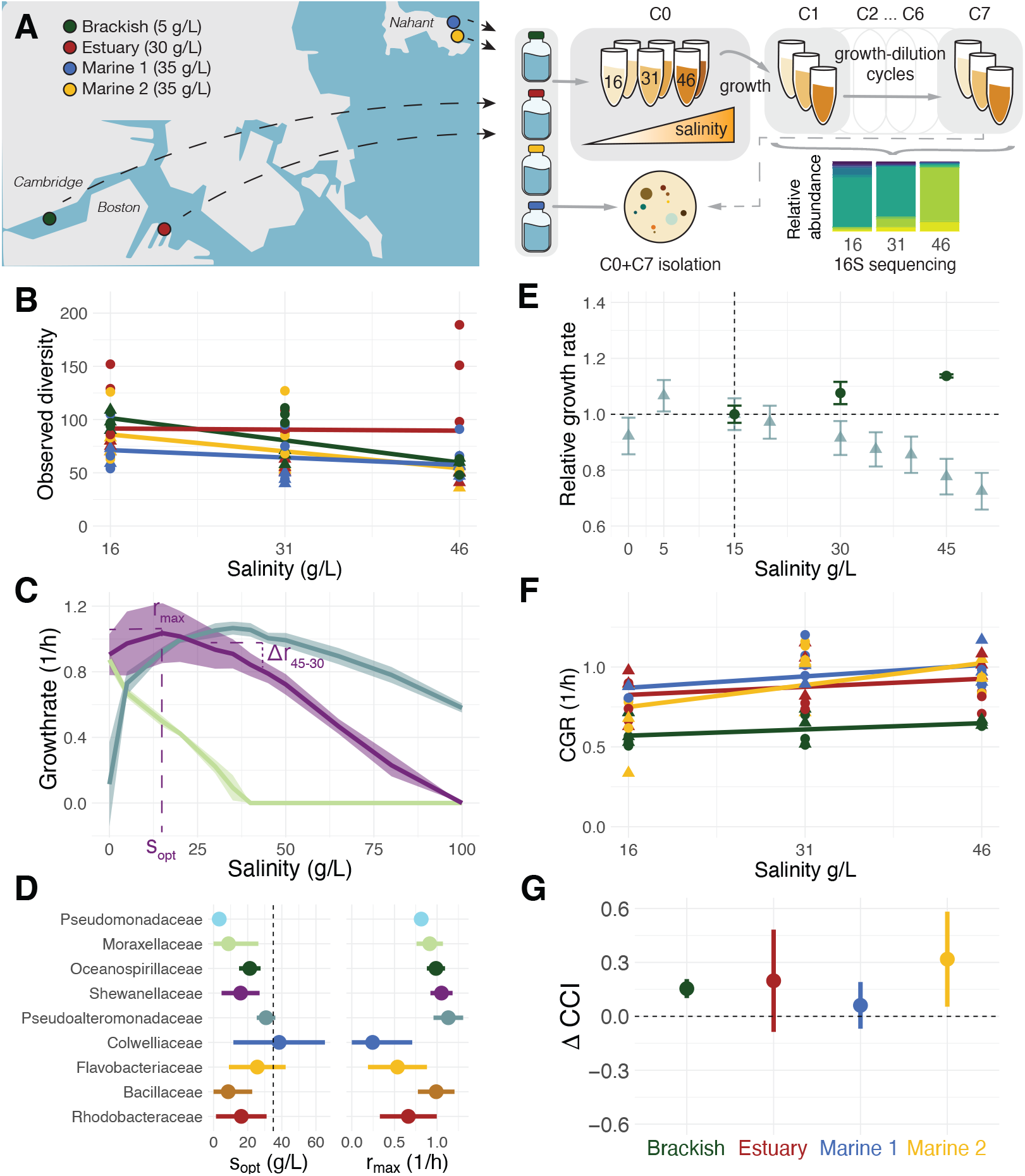
The community growth rate of aquatic microbial communities is more robust to an increase in salinity than expected based on individual species growth rates, due to a shift in community composition towards faster growing species. **A)** Schematic overview of the serial dilution experiment: natural aquatic communities from four locations were propagated over 7 48-hour cycles (C1-C7) at 3 salinities and 2 temperatures. Richness (observed number of ASVs) of the propagated communities at the end of the experiment (C6, *n* = 3 at 15 *◦*C and 20 *◦*C, denoted by triangles and circles respectively). Measured ‘salinity performance curves’ for three isolates (*Shewanella* in purple, *Acinetobacter* in lightgreen, *Pseudoalteromonas* in blue). The maximum growth rate (1/h) was extracted from 48hr OD600nm measurements at 12 salinities (n*>*3 replicates). **D)** Optimal salinity (*s*_*opt*_) and maximum growth rate (*r*_*max*_) for all families with at least 3 isolates. Drawn are mean *±* sd across n = 3 (Pseudomonadaceae) to n = 20 (Pseudoalteromonadaceae) isolates. In the case of *s*_*opt*_, the vertical dashed line indicates ocean salinity at 35 g/L. **E)** The community growth rate (green circles) and individual growth rates of isolates (light grey triangles) from the Brackish community. All values are scaled relative to the mean at 15 g/L. **F)** Community growth rate (CGR) for communities at the end of the experiment (C6, *n* = 3 at 15 *◦*C and 20 *◦*C, denoted by triangles and circles respectively). **G)** Change in the community composition index (CCI) between communities propagated at 15 g/L and 45 g/L (mean *±* se).

To understand the effects of salinity on the microbial community as a whole, we first characterized its effects on growth of the constituent species. We isolated 140 bacterial strains from the beginning (C0) and end (C7) of the serial dilution experiment, creating a library that includes *>* 60 microbial species, across 31 genera and 17 families. This library spans the known diversity of marine bacteria (Table S1, Fig. S7). Supplementing this with eleven isolates previously obtained at our marine sampling location [26, 27], we covered on average 95 *±* 6% of the observed community diversity at the end of the serial dilution experiment (at 97% 16S identity; Fig. S8). Maximizing phylogenetic diversity, we picked 85 isolates and measured their *salinity performance curve*, i.e. the effect of salinity (0-100 g/L sea salts) on maximum per capita growth rate (examples in Fig. 1C, all performance curves in Fig. S9). We find that isolates of the same family followed a similar salinity performance curve (Figs. 1D, S7, S9). Isolates from saline habitats obtained their maximal growth at higher salinities than isolates from the brackish isolation environment (Fig. S10). The curves typically reach maximum growth rate *r*^*max*^ at an optimal salinity *s*^*opt*^ between 0-35 g/L sea salts (ocean salinity is at 35 g/L), showing monotonically decreasing growth rates at salinities above this optimum (Fig. S9). Between 30 and 45 g/L, the growth rate of all isolates decreased by −0.06 *±* 0.01 1/h on average. Carrying capacities were less strongly impacted by increasing salinity than the growth rates (Fig. S6).

Next, we asked how the growth rates of individual bacteria relate to the growth rate of the community. The community growth rate governs the community’s recovery after perturbation and is intimately linked to community productivity, both crucial read-outs of community function. To compute the mean *community growth rate* (CGR), we weighed the growth rate of a species at a particular salinity by its abundance in the community propagated at that salinity (see Methods). The result is particularly striking for the Brackish community propagated at different salinities: while on average the growth rates of individual isolates from this community decrease above 5 g/L, the community growth rate actually increases between communities propagated at 15, 30, or 45 g/L (Fig. 1E). The same holds for the three communities isolated from more saline environments (albeit shifted to higher salinities): while the individual species growth rates decreased above 30 g/L, we find no sign of decreasing CGR with increasing salinity (Fig. 1F). If anything, the CGR even increased for some communities at higher salinity. The CGR is thus more robust to increased environmental stress than the growth of individual species.

To test whether the robustness of the CGR can be explained by a shift in community composition, we computed a quantity that describes the relative abundance of fast growing species. To obtain this *community composition index* (CCI) we assigned each species a single value which describes its relative growth rate compared to other species, here the species max growth rate at 30 g/L *r*(30), and computed the abundance weighted average of these species growth values for each community. We find that the CCI increases with increasing salinity for all starting communities (Fig. 1G). Increased environmental stress thus leads to a shift in species’ relative abundances in favor of faster growing species, which confers robustness to the mean community growth rate.

### Stressful environments enrich for faster growing species

To explain the observed enrichment of faster growing species in stressful environments, we developed a Lotka-Volterra model describing the ecological dynamics of microbial communities. We assume that bacterial species *N*_*i*_ grow at a species- and environment-specific growth rate *r*_*i*_(*s*), compete with each other with competition coefficients *a*_*ij*_, and are removed from the environment at a rate 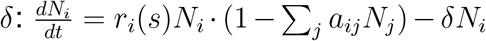 (Fig. 2A; Supplemental section S1). Continuous removal is common in natural environments, whether due to generalized predation or dilution [5, 6], and also a good approximation of the daily dilution process we employed in our experiments (supplementary note 2 in [6]). As a result of this mortality, the outcome of competition between two species depends on the removal rate *δ* and the growth rates of both species [4, 5, 6] (Supplemental section S1). Environmental stress, which affects the growth rates of all species, will thus affect the outcome of interspecies competition.

**Figure 2:**
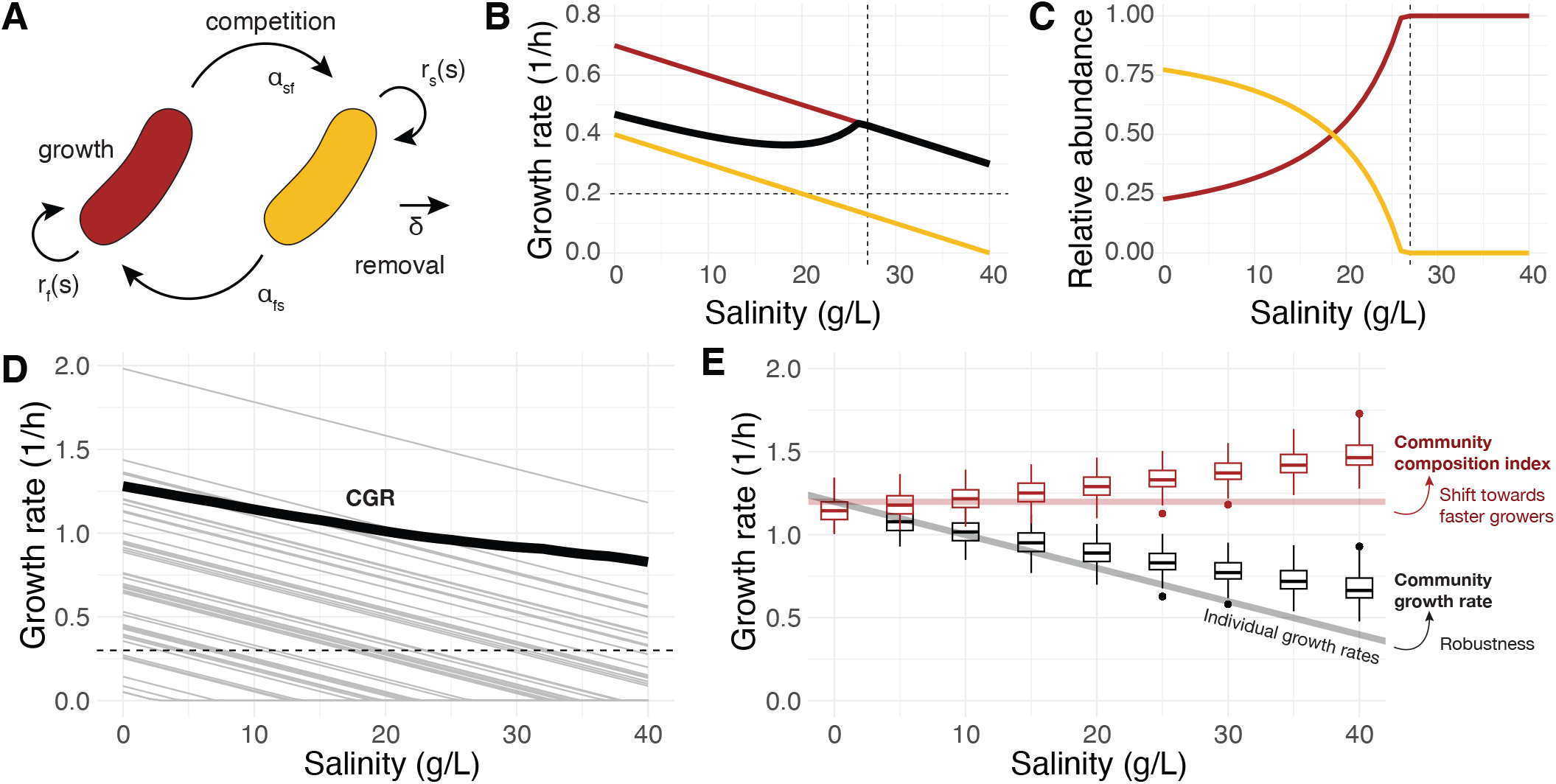
Modeling predicts that an increase in salinity shifts community composition towards faster growing species and confers robustness to the community growth rate. A) Model schematic of two species growing at rates *r*_*f/s*_(*s*), competing with inter-species interaction coefficients *α*_*fs*_, *α*_*sf*_, in the presence of removal from the environment at rate *δ*. B) Growth rates of the two competing species are assumed to decline at the same rate (*b* = −0.02; thin colored lines). The community growth rate (thick black line) declines more slowly. The dashed horizontal line indicates the strength of the removal rate (*δ* = 0.2). The dashed vertical line indicates the salinity at which the faster growing species first excludes the slower grower. C) Relative abundances of the two species at steady state at different salinities, with growth rates according to panel B. D) In a 50-species model where all growth rates decline at the same rate (thin grey lines), the community growth rate (thick black line) declines more slowly with increasing salinity. E) The robustness of the community growth rate is reproducible across 50 simulations of the 50-species communities (black boxplots). It is the result of a shift in community composition towards faster growing species (community composition index; red boxplots). If the community composition did not change with salinity (red line), the CGR would decline as the individual species growth rates (black line).

Having measured the functional relationship between growth rate and salinity, we can simulate how community composition will change as the environment deteriorates. Our salinity performance curve measurements typically show a salinity 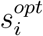 at which the maximal growth rate 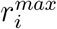 is reached, followed by an approximately linear decline in growth rate as salinity increases (here with slope *b*), until growth reaches 0 at 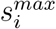 (Figs. 1C, S9, S11). When simulating a 2-species community, we see that the relative abundance of the faster growing species increases with increasing salinity (Fig. 2B/C). Notably, this shift in community composition confers robustness to the *community growth rate* (CGR; black line in Fig. 2B). The CGR declines more slowly than the growth rate of the species that make up the community.

This robustness is recapitulated in more complex simulated communities (*n* = 50), where the community growth rate increasingly approaches the salinity performance curves representing the fastest growing species (Fig. 2D). To describe the community composition index (CCI), we associate each species with its growth rate at 0 g/L salinity 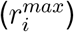 and compute the abundance weighted mean of these values for each of 50 simulated communities at a salinity *s*. Similar to the relative abundance of the faster grower in the 2-species case, the CCI increases with increasing salinity (red boxplots, Fig. 2E) and the corresponding CGR is more robust than the decline of the individual species growth rates (black boxplots and line, Fig. 2E). Importantly, this result only requires that the growth rate of all species declines with increasing salinity. We did not assume any effect of salinity on the interaction between species (*α*_*ij*_). As such, the compositional shift towards fast growing species and the resulting robustness of the community growth rate are expected to hold for any environmental stress that reduces the growth rate of most species.

The enrichment of faster growing species at higher salinities and the associated robustness of the CGR are not affected by model variations. A discrete model that more closely mimics the daily dilution in our experiments leads to the same qualitative outcomes (Fig. S12). As does variation in the relative slopes of the salinity performance curves between species (Fig. S13), or variation in the carrying capacities between species (Fig. S14). High removal rates *δ* and increasing slopes increase the initial strength of the CCI shift (Fig. S15, S16). They are made even more apparent in mixtures of bacteria with different optimal salinities (Fig. S17).

### Environmental stress can reverse pairwise competitive outcomes

The model suggests that the outcome of pairwise bacterial competitions will change in a predictable way as a function of salinity. We set out to test whether an increase in salinity indeed favors faster growing bacteria *in vitro*, using 8 pairs of isolates from our library, competing at four salinities (16, 31, 46, 61 g/L; Fig. 3A). For each pairwise competition, we started three replicate pairs at three different starting ratios (95:5, 50:50, 5:95), and propagated them every second day for 7 cycles at four different salt concentrations (16, 31, 46, 61 g/L). Across all conditions, the pairs reached stable final relative abundances (Fig. S18), with at least one salinity at which species coexisted at intermediate frequencies. The three different starting ratios led to similar final coexistence frequencies, revealing no evidence of bistability for any of the species pairs surveyed here.

**Figure 3:**
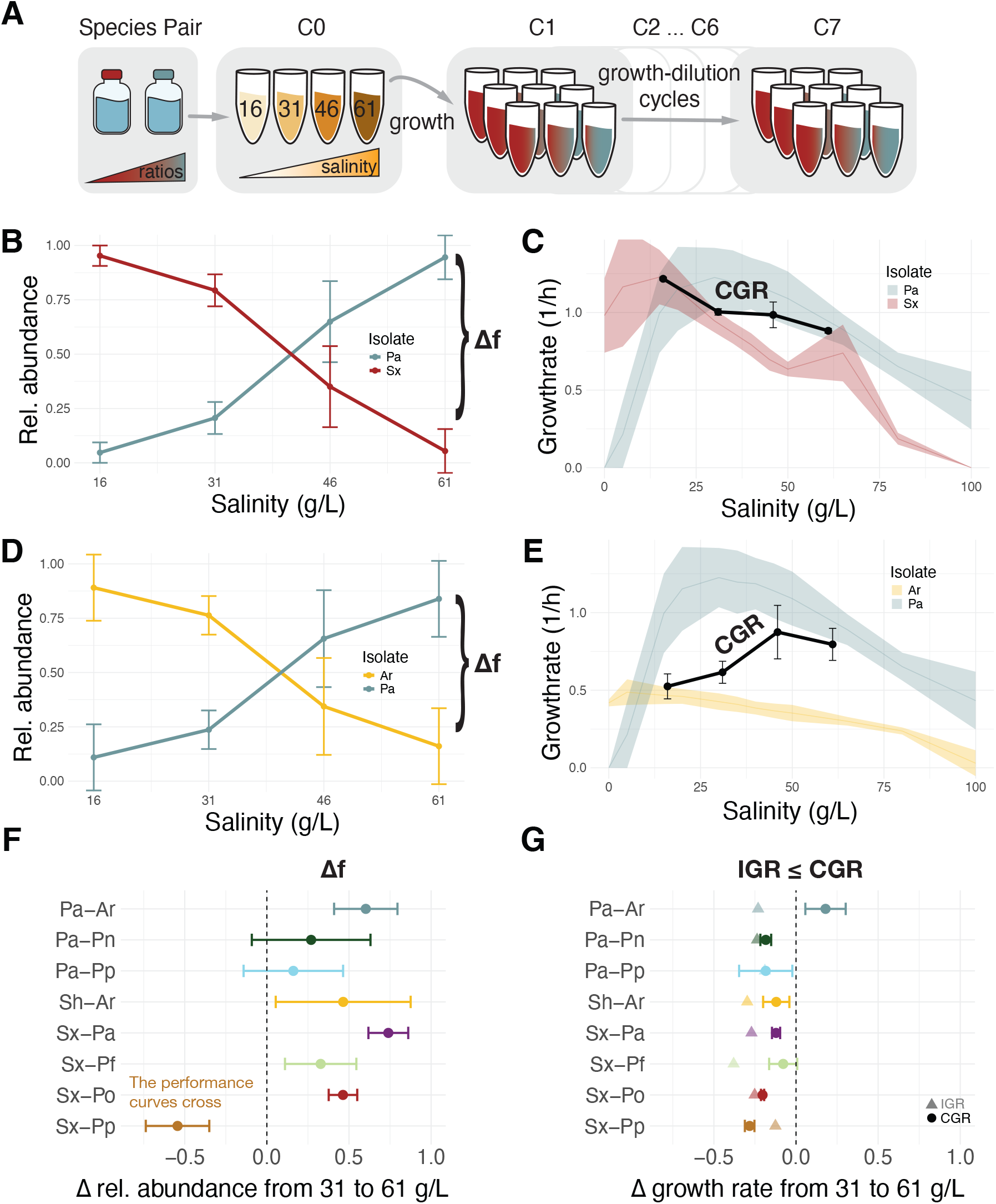
In pairwise co-culture, higher salinity favors the faster growing species. Experimental overview: we mixed three replicates of two species at three ratios (95:5, 50:50, 5:95) and propagated them at 4 salinities (16, 31, 46, 61 g/L) for 14 days (C1-C7). Relative abundance of *Pseudoalteromonas arctica* (blue; *Pa*) and *Shewanella xiamensis* (red; *Sx*) after propagation at 4 salinities (mean and sd across three biological replicates at three different initial ratios). **C)** Salinity performance curves for *Pa* and *Sx* (ribbons show mean *±* sd across *n* ≥ 3 replicates). The black line indicates the CGR of the pairs after 14 days of propagation at 16, 31, 46, 61 g/L salinity. **D, E)** As panel B, C but for the competition between *Pseudoalteromonas arctica* (blue; *Pa*) and *Albirhodobacter sp*. (yellow; *Ar*). **F)** The difference in the relative abundance of the faster growing species upon propagation at 31 and 61 g/L, across 8 species pairs. **G)** The difference in the community growth rate (CGR, circles) and mean isolate growth rate (IGR, triangles) upon propagation at 31 and 61 g/L, across 8 species pairs.

We found that, in all 8 competitions tested, the faster growing strain had an increasing competitive advantage at higher salinity (Fig. 3F, S18), and community growth rate declined less rapidly than the individual species growth rates (Fig. 3G). For example, in the competition between *Pseudoalteromonas arctica* (*Pa*) and *Shewanella xiamensis* (*Sx*) (Fig. 3B/C), the species coexisted at all salinities above 16 g/L, but the faster grower *Pa* became more abundant as salinity increases (Fig. 3B). As a result, the CGR stayed constant over a range of 30 g/L, while the growth rate of individual species declined, with *Sx* ‘s growth rate nearly halving across this range (Fig. 3C). This echos the findings of increasing CCI and robust CGR at higher salinity in our experiment with natural communities (Fig. 1), as predicted by our model (Fig. 2).

Strikingly, this dynamic is also observed in pairwise competitions where the slower growing species won at low salinities. When the same *Pa* was competed against an *Albirhodobacter sp*. (*Ar*), the latter won at low salinities (16 and 31 g/L) despite a substantial growth disadvantage (Fig. 3D/E). However, with increasing salinity (46 or 61 g/L) the faster growing *Pa* took over the community (Fig. 3D). In this case the CGR was not just robust to increased environmental stress, but actually increased with increasing salinity (Fig. 3E). We therefore observed that a stress-induced shift in community composition towards faster growing species lends robustness to the mean community growth rate, both in laboratory experiments with complex natural communities and in well-controlled species pairs.

### Environmental communities are enriched in faster growing species at higher salinity

Having gained insight into the effect of salinity on community dynamics *in vitro* and *in silico*, we next asked whether the increased abundance of faster growing species at high salinity can also be observed in natural microbial communities. In estuaries and inland seas, aquatic communities can be exposed to large seasonal or spatial salinity gradients. We identified six datasets of microbial communities sampled across such an environmental gradient: a 3-year time series taken at the Pivers Island Coastal Observatory (PICO; 26-38 g/L) [17], and spatial gradients in Chesapeake bay (1-20 g/L and 2-24 g/L) [18, 20], the Baltic sea (2-35 g/L) [16], the northern Gulf of Mexico along the Louisiana coast (0-26 g/L) [21], and lagoons along the Beaufort sea coast [19] (0-42 g/L; Fig. 4A; Table S2). Two of these datasets (the Louisiana coast [21], and the Chesapeake bay dataset of Cram *et al*. [18]) split the communities into different size fractions, allowing us to differentiate between free-living or particle-attached bacteria.

**Figure 4:**
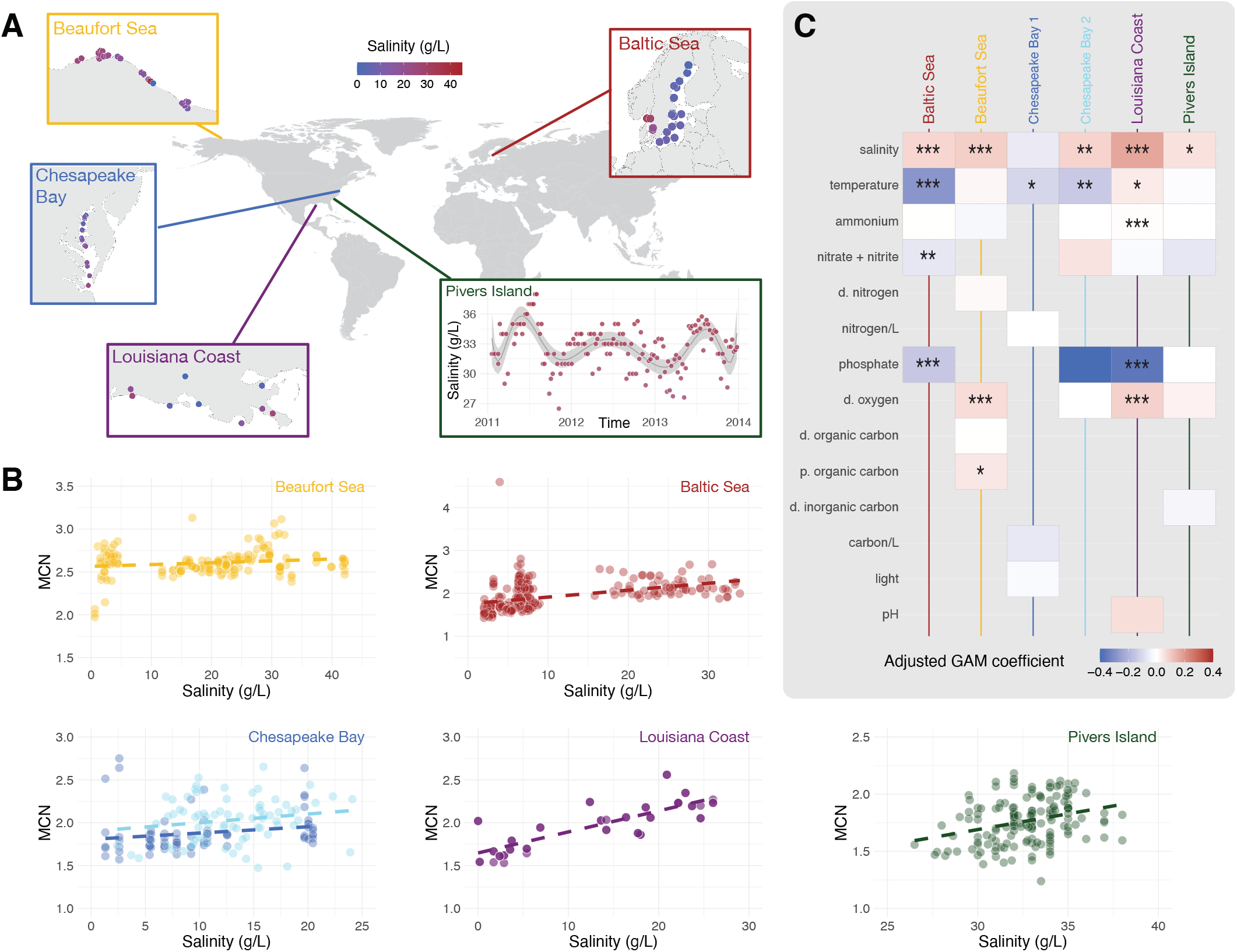
Higher salinity favors faster growers in natural estuarine communities, despite temperature and nutrient differences. A) Map indicating the sampling locations of the six datasets: Beaufort sea (yellow), Pivers island (green), Chesapeake bay (2x; dark blue corresponds to Cram *et al*. [18], light blue to Wang *et al*. [20]), the Louisiana coast (purple), and the Baltic sea (red). B) Mean copy number of the 16S rRNA gene (MCN) of each community sampled in the different datasets, plotted as a function of salinity at the location and time of sampling. C) Effect sizes of the different predictors of mean copy number. The color corresponds to the estimated parametric coefficient in the model, multiplied by the standard deviation of the environmental predictor (values below -0.4 were assigned the same color). Environmental predictors were abbreviated with “p.” for particulate, and “d.” for dissolved. The asterisks indicate significance at \*\*\**p <* 0.001, \*\**p <* 0.01, \**p <* 0.05.

Using genomic proxies to estimate the community composition index, we found an increasing proportion of fast-growing species at higher salinity for all datasets (Fig. 4B). We make use of the observation that the 16S rRNA operon copy number is a useful genomic proxy for the maximum growth rate a bacterium can attain [28, 29]. We verified this for our isolate dataset, and find a good correlation between 16S rRNA operon copy number and maximum growth rate at a species’ isolation salinity (Fig. S19). Mapping each community member to its estimated copy number [30] allows the calculation of an abundance weighted mean copy number (MCN) for the whole community. The MCN is commonly used to study microbial growth traits at the community-level, such as during microbial succession [31], or in response to increasing nutrients [32]. Here, we note that the MCN is the genomics-based equivalent of the community composition index (CCI) we calculated for the experimental communities [5]. We found that the MCN significantly increased as a function of salinity for the Baltic sea, Beaufort sea, Louisiana coast, and Pivers island datasets. While not significant, the Chesapeake bay dataset of Wang *et al*. also showed an increasing MCN with increasing salinity. In this case we treated different size fractions of the same water sample as independent communities for the Louisiana coast [21] and the Chesapeake bay dataset of Cram *et al*. [18]. However, our predictions best apply to free-living communities, and indeed we found that increasing salinity was associated with increased MCN in both datasets when considering only the free-living bacterial fraction (Fig. S20).

The increase in MCN with increasing salinity continued to be highly significant when accounting for all other environmental predictors measured in these datasets (Fig. 4B, Supp. Table S3). In fact, accounting for the effect of temperature additionally rendered the impact of salinity on MCN significant in the Chesapeake Bay dataset of Wang *et al*. Besides salinity, temperature most often had a significant impact on MCN (4/6 datasets), with evidence for increasing temperature favoring reduced MCN in 3/4. This has previously been described for other datasets of marine microbial communities and is consistent with our modeling framework, given temperature’s known role in increasing bacterial growth rates [4, 5]. In addition, we found that increasing phosphate was associated with a significantly reduced MCN in two datasets (the Baltic sea and Louisiana coast), while dissolved oxygen was associated with a significantly increased MCN (Beaufort sea and Louisiana coast). To conclude, the community shift towards faster growing species at high salinity that we observed *in vitro* is recapitulated in environmental datasets despite differences in temperature and nutrient availability.

## Discussion

In this work, we set out to investigate how environmental stress impacts community composition and function. We showed that microbial community growth rate – which sets the pace for fundamental ecosystem functions, including biomass production and the ability to recover from perturbations – is remarkably robust to environmental stresses that reduce species’ growth rates. This robustness is the result of a change in community composition towards faster growing species. We have demonstrated this in the context of natural aquatic communities propagated at different salinities, as well as pairwise competitions and environmental estuarine datasets.

This compositional shift and community robustness likely extends beyond salinity to other stressors, including pH, temperature and chemical pollutants. Our modeling framework lacks environment- or stressor-specific assumptions and would make similar predictions for any stressor that reduces the growth of most bacteria. Indeed, prior research noted that growth of a mixed community of bacteria proved more robust to the influence of chemical pollutants than growth of the constituent members [33]. In pairwise competitions, faster growth proved the most important determinant to predict competitive outcomes in the presence of increasing sublethal concentrations of antibiotics [34], increasing daily dilution rates [6], and reduced temperature [4]. For temperature this effect was also observed in environmental datasets, where a decrease in temperature corresponds to a decrease in the growth rate of species and leads to the same community shift as an increase in salinity [5]. All these observations would be consistent with emergent robustness of the community growth rate due to a compositional shift towards faster growers in the presence of environmental stress.

Environmental stress is often characterized by changes in not only one, but multiple environmental variables. Here, we found that the impact of salinity on the composition of natural communities was robust to environmental confounders such as temperature and nutrient concentrations. However, interactions between changing variables may still structure bacterial growth in non-trivial ways. For instance, higher temperatures can expand the range of salinity tolerance for some species ([35] and own preliminary data). Similarly, there could be an interesting interplay between nutrient limitation and salinity stress. If nutrient stress acts synergistically with salinity in reducing growth rates, we would expect joint exposure to both stresses to select for fast growing species even more strongly than salinity alone. Yet, slow-growing, oligotrophic, bacteria are often considered specialized for growth at low nutrient concentrations. This suggests their growth rates may not decrease as fast as for other species at low nutrient concentrations, leading them to become the de-facto faster growers. Such scenarios where the environmental variable impacts some species more than others, or changes their level of competitive interactions, will cause the simple predictions from our model to become more complex. More growth measurements are needed to parametrize performance functions across multiple simultaneously varying variables, and to assess the predictability of community-level behavior under multivariate stressors.

Importantly, while environmental stress did not affect community growth, it did lower community diversity. This may negatively impact a community’s ability to perform more complex functions, such as nitrification or bioremediation, if those functions are restricted to impacted taxa [7, 36]. This highlights a dilemma in how to assess ecosystem health and functioning: high community biomass and growth rates might be achieved even when community diversity is already severely impacted. The loss of genetic diversity at both within- and between species levels is irreversible without influx from the external species pool and may lead the community to enter a long-term alternative stable state. The loss of functional diversity can also reduce resilience to future perturbations with a different stressor. This underlines the importance of assessing diversity also in communities that seem robust to stress in terms of biomass production and/or growth rate.

To conclude, we have uncovered a general principle governing the response of communities to environmental stress. Increasing stress increases the relative importance of growth rates - rather than competitive abilities - in dictating community coexistence outcomes. As a result, increasing stress causes the community composition to shift towards faster growing species, which confers robustness to the mean community growth rate.

## Methods

### Sampling and C0 species isolation

We sampled water at three locations around Boston on March 31st 2023. The locations span a natural salinity gradient: The Charles River at the MIT sailing pavilion (“Brackish”; 4 g/L; 10 *◦*C), the Boston harbor near the Institute for Contemporary Art (“Estuary”; 30 g/L; 6 *◦*C), and the ocean at Canoe Beach, Nahant (“Marine 1”; 35 g/L; 7 *◦*C). Temperature and salinity were recorded at the time of sampling (refractometer kindly borrowed from the Fakhri lab, MIT). Per location, we filtered 2L of water using a 63 *µ*m filter (kindly borrowed from the Cordero lab, MIT; filter pre-washed with water from the respective location) to remove particulate matter and larger eukaryotic cells. We concentrated the samples by centrifuging them at 4000 rpm for 5 minutes and keeping only the bottom 10% of the sample. This resulted in 100 mL of concentrated water per inoculum sample.

We additionally sampled brown macroalgae (likely *Ascophyllum nodosum*) at Nahant. To obtain the algae-attached communities, we placed seaweed blades into 8 50 mL conical tubes (Falcon) and vortexed them in seawater from Nahant for 2 minutes prior to sieving with the 63 *µ*m filter. The 8 tubes were centrifuged at 4000 rpm for 5 minutes, we combined the bottom 10% (40 mL) into a single falcon tube. The resulting concentrated water was used as fourth aquatic community sample (‘Marine 2’).

The majority of the sample was used to inoculate a serial dilution experiment (described below). The rest was frozen both directly and with 30% glycerol at -80 *◦*C. To isolate species from the original communities, we plated 150 *µ*L concentrated inoculum onto 2 replicate 2% marine broth agar plates (MB; Becton Dickinson; bacteriological agar VWR). These were left to grow at room temperature for 3 days. Then, we visually inspected the plates, picked colonies, and streaked them out individually (‘C0 isolates’). We revisited plates after 7 days, and picked 8 additional colonies in the same manner. For permanent storage, we used the streaked plates to pick one colony per isolate and transferred it into a 1mL deepwell plate (Eppendorf) with 400 *µ*L MB. After 2 days of growth at room temperature on a shaker at 1350 rpm, we added 400 *µ*L of sterilized 50/50 glycerol to each well, and transferred the plates to the -80 *◦*C freezer.

### Serial dilution experiment and C7 species isolation

To obtain stable communities at different salt conditions, we ran a 14-day serial dilution experiment in which the 4 inocula were adapted to 3 different salt concentrations (16, 31, 46 g/L sea salts). We used undiluted marine broth (MB, Becton Dickinson) as base medium (37 g/L MB corresponds to 31 g/L sea salts). With ‘sea salts’ we denote the 5 primary salts present in MB, at their respective relative proportions: 19.45 g/L Sodium Chloride, 5.9 g/L Magnesium Chloride, 3.24 g/L Magnesium Sulfate, 1.8 g/L Calcium Chloride, 0.55 g/L Potassium Chloride. To obtain the other salt concentrations, the amount of nutrients was kept fixed: the lower salinity (16 g/L) was reached by diluting the MB and adding nutrients (peptone, yeast, ferric citrate), and the higher salinity (46 g/L) by supplementing MB with sea salts. All three concentrations of MB were filter sterilized by passing through a.2 *µ*m filter (Stericup, Millipore) and stored in the dark at 4 *◦*C.

To start the experiment, we inoculated 3 replicates of each aquatic sample (10 *µ*L concentrated inoculum) into a 96 well 500 *µ*L deepwell plate (Eppendorf) with 290 *µ*L MB per well, at 3 different salt concentrations (16, 31, 46 g/L sea salts). This yielded a total of 9 inoculated communities for each of the 4 samples, the remaining 60 wells were used as spacing and blanks to control for contamination. We incubated two separate plates at 15 and 20 *◦*C. Plates were kept on benchtop shakers (Titramax 100) operated at 1200 rpm. Every second day for 2 weeks, we diluted the culture in each well 1:30 and transferred the communities to a plate with fresh culture medium (using the Integra VIAFLO96). After every transfer, we used 100 *µ*L from the old plate to measure pH (Thermo Fisher Orion Star A211) and OD600 (Tecan Infinite M Nano). The remainder of the old plate (190 *µ*L per well) was then stored at -80 *◦*C, to be used for DNA extraction.

After 7 dilution cycles (14 days; the end of the experiment), we diluted each community 10^−7^ in PBS. We plated 50 *µ*L onto 150×15 mm petri dishes with 2% marine broth agar (MB; Becton Dickinson; bacteriological agar VWR). The three replicates of the Marine 1 inoculum at 16 g/L salt at 15 *◦*C and 20 *◦*C were additionally plated at a dilution of 10^−5^. We counted all colonies on these plates, picked representative colonies displaying different morphology, and individually streaked them onto 100×15 mm 2% MB agar plates (‘C7 isolates’). Single isolates were picked from these plates, grown in 1 mL MB overnight, combined with 0.5 mL sterile glycerol and stored at -80 *◦*C. Additionally, at dilution cycle 7, we transferred the deepwell plates one more time to allow for long-term storage of the full communities. After pH and OD600 measurement on day 16, we added 200 *µ*L 50/50 glycerol to these cycle 8 plates and stored them at -80 *◦*C.

### Isolate 16S rRNA sequencing and species identification

C0 and C7 isolates were streaked onto fresh 2% MB agar plates from the -80 *◦*C stock and sent for 16S rRNA gene Sanger sequencing with Azenta Life Sciences. We used their inhouse pipeline to trim and merge the forward and reverse reads. We assigned taxonomy using DADA2 [37] and the SILVA database (v138.1) [38].

We aligned the 16S genes with MAFFT [39], and used RaxML [40] with a GTR model with Γ site variation to estimate a maximum likelihood phylogeny for all isolates.

### Community 16S rRNA sequencing and ASV calling

To obtain community 16S sequencing data, we sent each replicate community from the C1, C3, C5, C6, C7 cycles, as well as two replicates per C0 inoculum for sequencing (Novogene). We used v4-v5 primers to obtain 250bp paired-end reads on an Illumina NovoSeq 6000. In total, we sequenced 368 communities.

We used the DADA2 [37] pipeline to trim and filter the sequencing reads, infer amplicon sequence variants (ASVs), merge paired reads, and remove chimeras. We additionally used it to assign species taxonomy according to the SILVA database (v138.1) [38]. For further analyses we removed spurious ASVs that were classified as mitochondria or non-bacterial, and ASVs that were only found in a single sample.

We calculated the species richness by counting the number of different ASVs in a sample. We use rrnDB to assign each ASV a copy number. We assigned each ASV a copy number according to the lowest taxonomic rank for which a value was available in the database. Using these copy numbers, we calculated the true relative abundance of each ASV. For each sample we additionally calculated the mean copy number (MCN) which denotes the average copy number of the ASVs in the sample (weighted by their relative abundance). ASVs were matched to isolate 16S rRNA gene sequences using BLAST at *>* 97% identity [41].

Four samples from C7 showed unexpectedly high diversity (all three replicates of the estuary community propagated at 46 g/L and 15*◦*C, as well as replicate R1 of the marine 1 community propagated at 46 g/L and 15*◦*C). These were all co-located with C0 samples in the first two rows of one of the plates sent for sequencing, which leads us to believe there was a contamination during transport or sample handling. As a result, we used the community data from cycle C6 for all subsequent analysis.

To understand changes to community composition over time, we computed pairwise Bray-Curtis distances between communities of the same replicate condition, sequenced on following days in the serial dilution protocol. All distances were computed using the vegdist function from the vegan package [42].

### Growth rates

To measure the growth of different aquatic isolates across a gradient of salinity, we used 12 different media with the nutrients of normal MB (except Sodium Fluoride) and varying amounts of the 5 primary sea salts (NaCl, MgCl, MgSO, CaCl, KCl) in the same ratio as in MB. A MB-nutrient only solution was combined with MB with 100 g/L salts in different ratios to obtain 0, 5, 15, 20, 30, 35, 40, 45, 50, 65, 80, 100 g/L MB solutions. The two stock solutions were filter sterilized by passing through a.2 *µ*m filter (Stericup, Millipore) and stored in the dark at 4*◦*C.

Strains were streaked from -80 *◦*C stocks onto 2% MB agar, grown at room temperature for 2-4 days and stored at 4 *◦*C. For each strain, we picked a replicate isolate on three separate days, and grew it in 3 mL MB in a 17×100 mm culture tube (VWR) for 48 hrs at 20 *◦*C with spinning (∼ 100*±*25 rpm) on a rotary lab suspension mixer (TMO-1700, MRC Lab).

We diluted these overnight cultures 100-fold in MB, and used them to inoculate a 96 well plate (hereafter called the ‘measurement plate’). We added 2 *µ*L diluted isolate culture to wells containing 200 *µ*L MB at 12 different salinities. The sides were taped against evaporation. Growth rates were measured on a Tecan Infinite M Nano, for 48 hours at 20 *◦*C.

We used the R package gcplyr (v1.11.0) [43] to estimate growth rates from the OD measurements. Specifically, we smoothed the measured OD across 21 minutes (7 datapoints; sliding window) and used gcplyr to estimate the derivative of the log OD curve. This derivative corresponds to the average growth rate across a 1-hr window. We denote the maximum growth rate for growth at this salinity *r*_*max*_. To exclude spurious results, we did not assess the derivative in the first 3 hours of growth and if the OD for a given strain at a given salinity did not exceed 0.1, we set *r*_*max*_ = 0.

We used the package *segmented* in R, to fit both a linear and a piecewise linear function with two segments to the measured salinity performance curves. If the adjusted R squared of the linear fit was better than that of the 2-segment model, the linear fit was kept and *s*^*opt*^ assumed to be 0. The latter (*s*^*opt*^ = 0) was also assumed if the 2-segment model inferred a negative slope for the first segment. In all other cases, *s*^*opt*^ was assumed to be at the inferred breakpoint of the two segments.

To determine the *community growth rate* (CGR), we calculated the abundance weighted mean of the realized growth rates of all species in the community. That is, if *f*_*i*_(*s*) denotes the measured relative abundance of strain *i* in a community at salinity *s* and 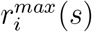 denotes. the maximum growth rate of that strain at that salinity, 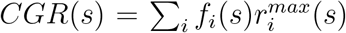. Instead, the *community composition index* (CCI) at a salinity *s* is defined as *CCI*(*s*) = ∑ *f*_*i*_(*s*)*r*_*i*_ Here the characteristic growth rate *r*_*i*_ of a species *i* can either be its maximum growth rate across all salinities measured (*r*^*max*^(*s*_*opt*_)) or the growth rate at a salinity that is ecologically relevant (e.g. the salinity of the isolation environment, *r*^*max*^(30*g/L*)).

### Pairwise Competitions

Using the C0 and C7 isolates, we selected 8 strains for pairwise competitions (*Pseudoalteromonas arctica Pa, Shewanella xiamensis Sx, Shewanella sp. Sh, Albirhodobacter sp. Ar, Pseudoalteromonas ostrae Po, Pseudoalteromonas nitrifaciens Pn, Pseudomonas fragi Pf*, and *Psychrobacter piscatorii Pp*). The isolates were selected based on 16S similarity to dominant ASVs at the end of the serial dilution experiment, discernible phenotypes on MB agar plates, and different salinity performance curves.

To reach the steady state outcome of the competition, we propagated the 8 pairs for 14 days at 4 different salt concentrations (16, 31, 46, 61 g/L sea salts) and 3 different starting ratios (95:5, 50:50, 5:95). The media was prepared as for the growth rate measurements.

To start the experiment, we grew 3 biological replicates of all 8 strains in MB for 48 hours, washed them twice in PBS, OD standardized to the lowest OD, mixed strains in correspondence with the intended starting ratio, and inoculated 10 *µ*L of this strain mixture into four (one per salinity) 96 well 500 *µ*L deepwell plates (Eppendorf) with 290 *µ*L growth medium per well. Monoculture growth controls were included for each biological replicate. Unfortunately the control for isolate 6 was contaminated (50-50) by isolate 8 during inoculation, and not used for further analysis. The four plates were incubated at room temperature (20-21 *◦*C) on a benchtop shaker (Titramax 100). Every second day, we diluted the culture in each well 1:30 and transferred the communities to a plate with fresh culture medium (using the Integra VIAFLO96). After every transfer, we used 100 *µ*L from the old plate to measure OD600 (Tecan Infinite M Nano).

At the start of the experiment and after cycles C1, C3, C5, C6, C7, we diluted each plate 10^−6^ and 10^−7^ in PBS, spot-plated 10 *µ*L per well, and counted the corresponding colonies after 2-3 days. For pairs *Pa*-*Sx, Sx* -*Pp, Sx* -*Pf, Sx* -*Po, Pa*-*Pp*, and *Pa*-*Pn* we used the counts at 10^−6^ for further analysis. For the two pairs with *Ar*, i.e. *Pa*-*Ar* and *Sh*-*Ar*, we used dilution 10^−7^.

### Modeling

We describe our modeling framework in more detail in Supplementary Section S1. In brief, we used a generalized Lotka-Volterra model with constant dilution rate *δ* to model the competition between two or more species, *N*_*i*_:

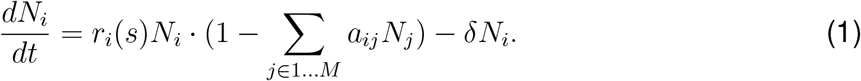

Here *r*_*i*_(*s*) denotes the growth rate at salinity *s*. The self-inhibition terms are all set to *a*_*ii*_ = 1 (equivalent to normalizing by carrying capacity), and the terms *a*_*ij*_ describe the effect of species *N*_*j*_ on the growth of *N*_*i*_.

We tested the effect of this normalization of the carrying capacity by comparing it to three other scenarios: one where the *a*_*ii*_ were sampled in the same way as the *a*_*ij*_ (‘free’), sampled as *a*_*ij*_ and then reordered such that the fastest growers have the highest *a*_*ii*_ (i.e. the lowest carrying capacity; ‘trade-off’), or sampled as *a*_*ij*_ and then reordered such that the fastest growers have the lowest *a*_*ii*_ (i.e. the highest carrying capacity; ‘trade-up’). We also simulated a version of this model with discrete dilution where the continuous removal rate *δ* = 0.2 was replaced by a 122x dilution every 24 time steps.

We simulated these equations deterministically using the R programming language [44]. Growth rates *r* were sampled from a normal distribution 𝒩 (0.7, 0.2), *a*_*ii*_ = 1, and *a*_*ij*_ were sampled from a uniform distribution on the interval (0, 2*α*). The mean interaction strength is *α* = 0.25 for the main-text figures.

To study the effect of changing salinity, three different scenarios were assumed for *r*_*i*_(*s*): a parallel decline where 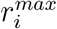 is sampled but the slope is fixed for all species, a converging decline where 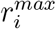 is sampled and 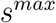 is fixed for all species, and a diverging decline where 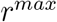 is fixed for all species but the slope is sampled (Supp. Fig. S13A).

### Environmental metagenomic samples

We searched Pubmed for published research papers that sampled aquatic microbial communities at different salinities and reported the 16S sequencing data. We added articles that were identified via independent routes or from forward/back citations of articles previously identified. By screening abstracts for relevance, we narrowed several hundreds of papers down to a set of roughly 20 articles, for which we read the full text to determine their suitability for inclusion in our comparison. Articles were excluded if the range of salinities measured spanned less than 10 g/L, involved non-aquatic systems, contained *<* 20 data points, contained no information on confounding environmental variables such as temperature, or if the 16S data was not available. This process resulted in the identification of four studies [16, 17, 18, 19]. Based on recommendation by an anonymous reviewer, we later added two additional independent datasets [32, 21].

For all studies we downloaded 16S sequencing data, metadata, and all measured environmental variables (Table S2). For four of these [18, 19, 20, 21] only raw reads were available, which required pre-processing with DADA2 to obtain the table of ASV abundances. We followed the same steps as detailed in methods section “Community 16S rRNA sequencing”, with trimming parameters optimized to each dataset.

We fitted general additive models to our data, taking into account all measured environmental variables for each dataset (provided they were not highly correlated with other included variables).

## Data availability

All code and data is available at: https://github.com/JSHuisman/salinity.

## Acknowledgements

The authors thank two anonymous reviewers for their comments which significantly strengthened this manuscript, in particular by drawing our attention to two additional environmental datasets. We further thank Shreyansh Umale and other members of the Gore lab for helpful discussions about this work. JSH was supported by Human Frontier Science Program (HFSP) Postdoctoral Fellowship LT0045/2023-L.

## Author contributions

JSH designed the study; performed sampling, in vitro experiments (serial dilution, growth and pairwise competition) and modeling; contributed to the environmental data analysis; visualized results; and wrote the first draft of the manuscript. MDB designed the study; performed sampling and environmental data analysis; contributed to serial dilution experiments and data visualization; and edited the manuscript. JG designed the study; supervised in vitro experiments and modeling; and edited the manuscript.

## S1 Mathematical models of competition

We used mathematical modeling to understand the impact of a deteriorating environment on the competition between two or more species. The generalized Lotka-Volterra model is a powerful framework to describe such ecological competition, summarizing complex biological processes (e.g. resource competition, competition for space or toxin production) into either self-inhibition (*a*_*ii*_) or the net effect of species *N*_*j*_ on the growth of *N*_*i*_ (*a*_*ij*_). In serial dilution experiments in the laboratory, as well as most natural environments, species will also be removed (or die) at a constant rate *δ*.

This yields the general model equations governing the population size *N*_*i*_ of species *i*:

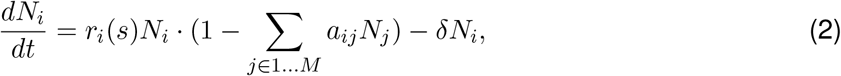

where *r*_*i*_(*s*) denotes the growth rate in environment *s*, and the self-inhibition terms are *a*_*ii*_ *>* 0. The interaction terms between both species *a*_*ij*_ can take any real number.

### S1.1 Pairwise Model

To study the dynamics of these equations in more detail, we first focus on the case with two species (*N*_1_, *N*_2_). *Dt*

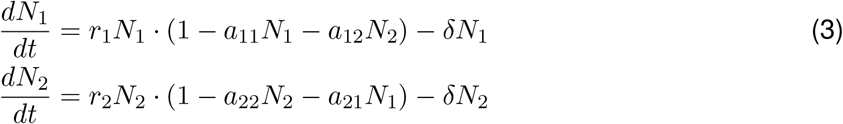

The classical form of 2-species LV equations can be obtained by integrating the dilution rate into new *effective* parameters [6, 4, 5]:

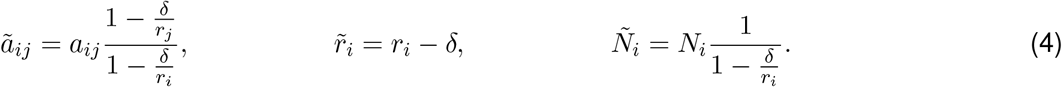

For the first ODE this yields:

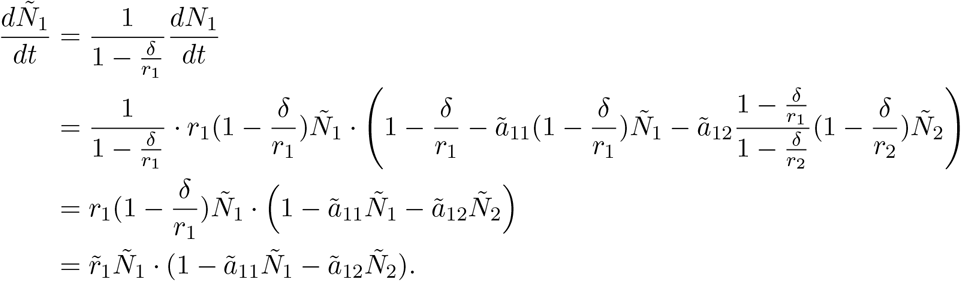

So we recover the new system of equations:

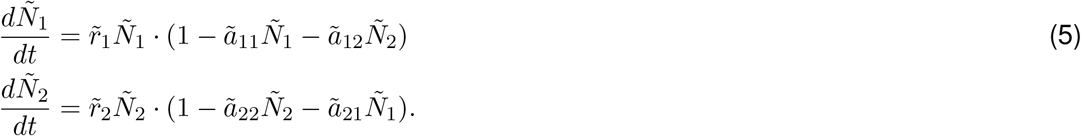

#### Steady state outcomes

To obtain an analytical expression for the steady-state outcome of the competition between *N*_1_ and *N*_2_, we solve ODE 3 at 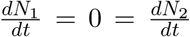. We find the following four outcomes:

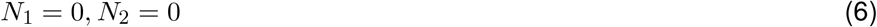

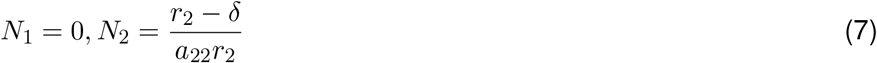

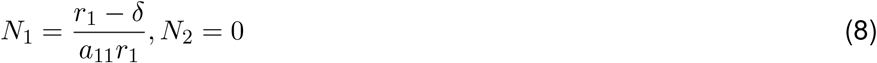

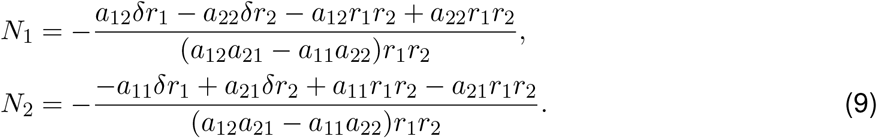

This is equivalent to:

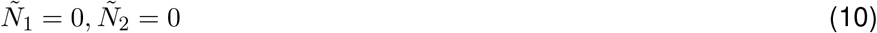

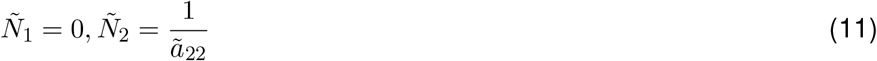

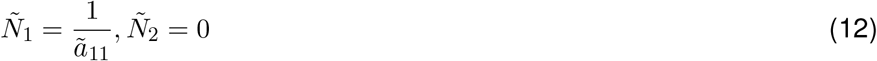

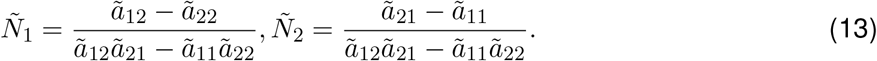

#### Coexistence, bistability or exclusion

The fourth steady state allows positive species abundances when *ã*_12_ *> ã*_22_ and *ã*_21_ *> ã*_11_ or *ã*_12_ *< ã*_22_ and *ã*_21_ *< ã*_11_. It is easy to see that the first set of constraints yields a bistable system, where species 1 more strongly inhibits species 2 than species 1 inhibits itself and vice versa, while the second set of constraints yields coexistence [45].

As such we find the following conditions for the canonical version of the LV model with effective parameters *ã*_12_, *ã*_21_:

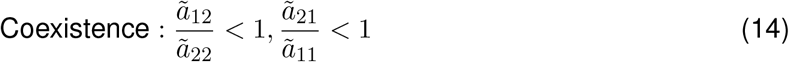

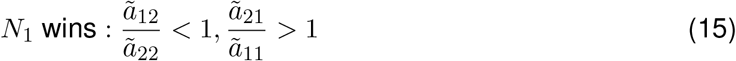

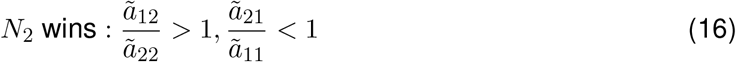

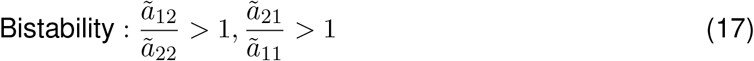

And for the corresponding equations with the full parameters:

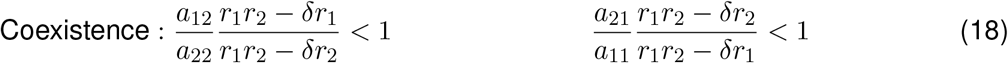

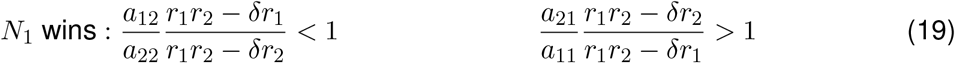

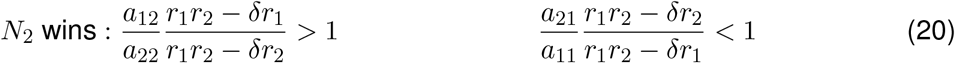

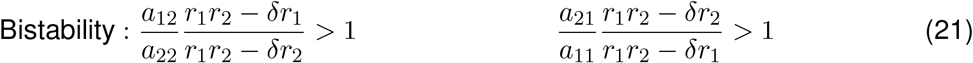

In the coexistence state (equations 9 or 13) the ratio of *N*_1_*/N*_2_ is given by:

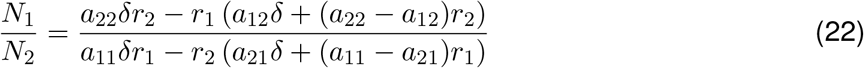

Or equivalently:

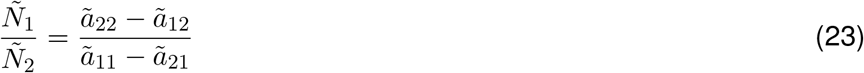

#### Changing salinity

The previous results hold generically, for any two species whose competition can be described in a Lotka-Volterra framework with dilution (eq. 3). However, now we can ask how these steady state outcomes change as the environment changes. We assume that the interspecies interactions (*a*_*ij*_) and carrying capacities (1*/a*_*ii*_) do not change with salinity. Then we see that for the three nontrivial steady state outcomes (eq. 9), the final steady state abundance of each population is determined by the functional relationship between growth rate and salinity. For simplicity, we assume specific scenarios:

- Parallel growth curves: the two species exhibit different maximal growth rates 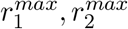, which decrease linearly at the same rate *b*:

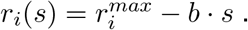
- Converging growth curves: the two species exhibit different maximal growth rates 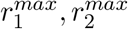, which decrease linearly to the same point *s*^*max*^:

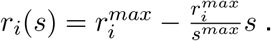
- Diverging growth curves: the two species exhibit the same maximal growth rate *r*^*max*^, which decrease linearly at different rates *b*_1_, *b*_2_:

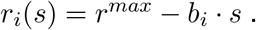

In each case, we want to know how the ratio between both species *N*_1_*/N*_2_ changes with increasing salinity. To do so, we calculate the derivative of eq. 22 with respect to *s* upon substitution of these different linear functions for *r*_1_(*s*), *r*_2_(*s*).

Parallel growth curves:

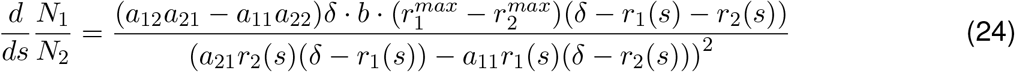

Under the conditions that yield coexistence, i.e. *a*_12_*a*_21_ *< a*_11_*a*_22_, and assuming that *r*_1_ *> r*_2_ and *δ < r*_1_ + *r*_2_, we find that 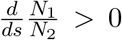. The proportion of the faster growing species increases with increasing salinity.

We can perform the same analysis for converging growth curves:

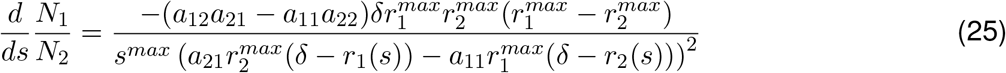

or diverging growth curves:

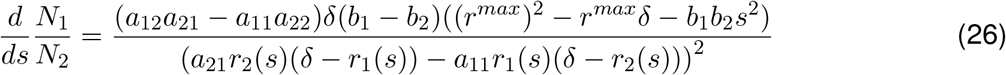

Across all scenarios a high dilution rate *δ* contributes to a strong effect of salinity on the proportion of the faster growing species.

## S2 Supplementary Figures

**Figure S1:**
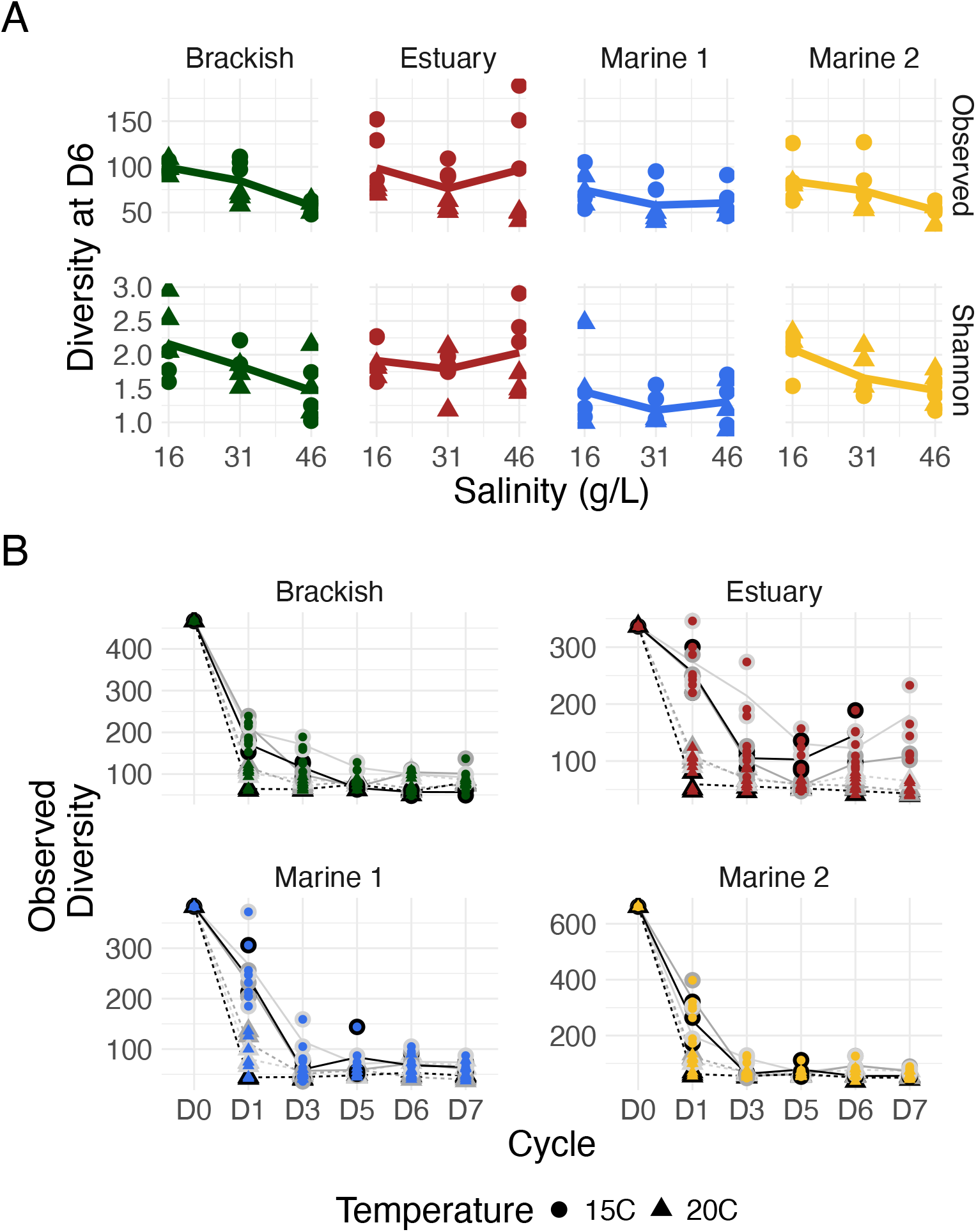
Diversity of the communities during the serial passaging experiment. A) Observed (top row) and Shannon diversity (bottom row) of the propagated communities after 6 cycles. Diversity metrics are based on the 16S community sequencing data, assuming each ASV is a unique species. Colors represent the different starting communities. B) Dynamics of the observed community diversity over time. Lines follow the mean of three replicate communities, split by salinity (16 g/L in lightgrey, 31 g/L in darkgrey, 46 g/L in black) and temperature (15C as solid, 20C as dashed lines) of propagation.

**Figure S2:**
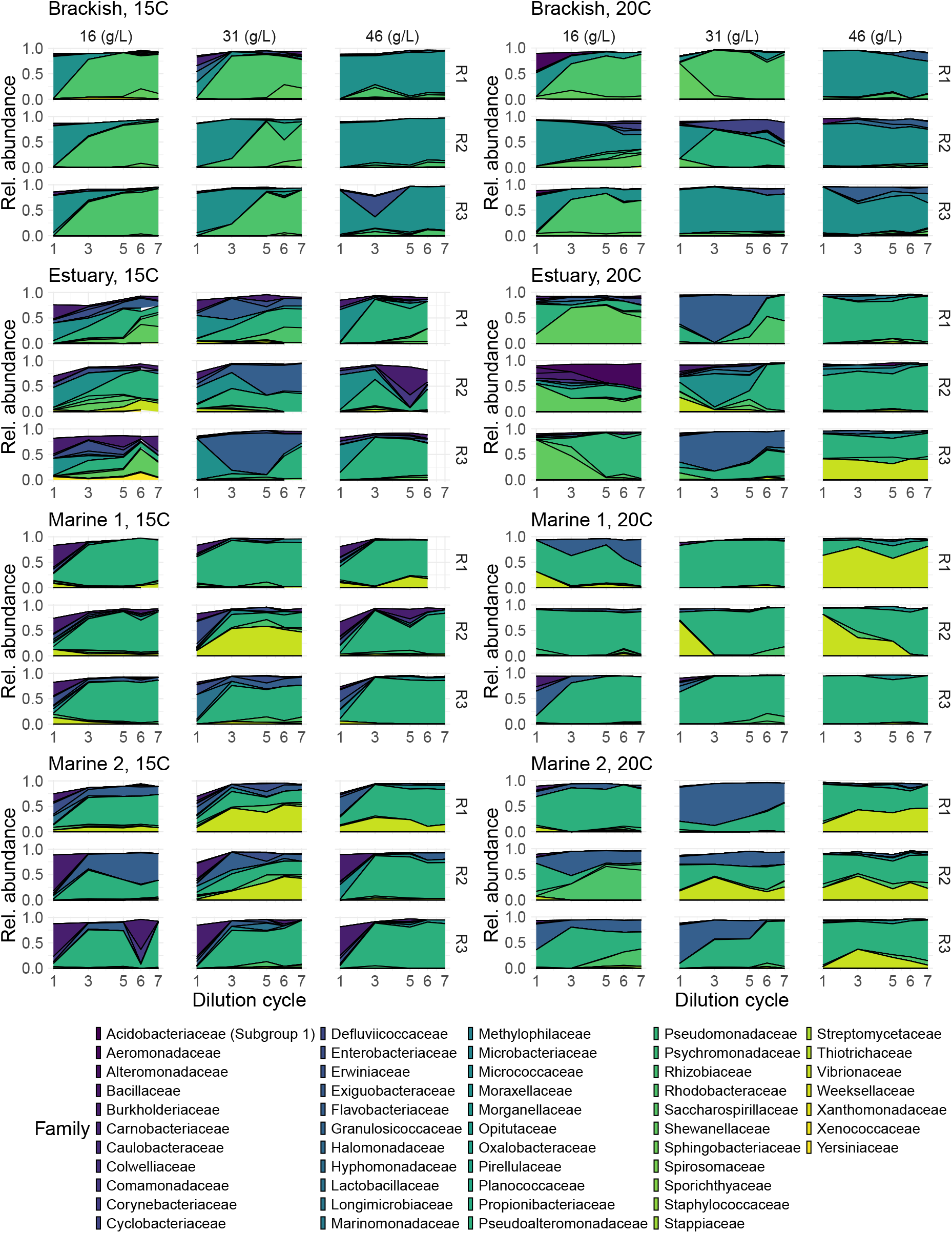
Community composition over 7 growth-dilution cycles. Communities are separated by the propagation temperature (columns; 15 or 20 *◦*C) and the source community (rows; Brackish, Estuary, Marine 1, Marine 2). Three replicate communities (R1-3) were propagated for each condition. ASVs are colored by taxonomic family, and only ASVs at greater than 1% relative abundance are depicted.

**Figure S3:**
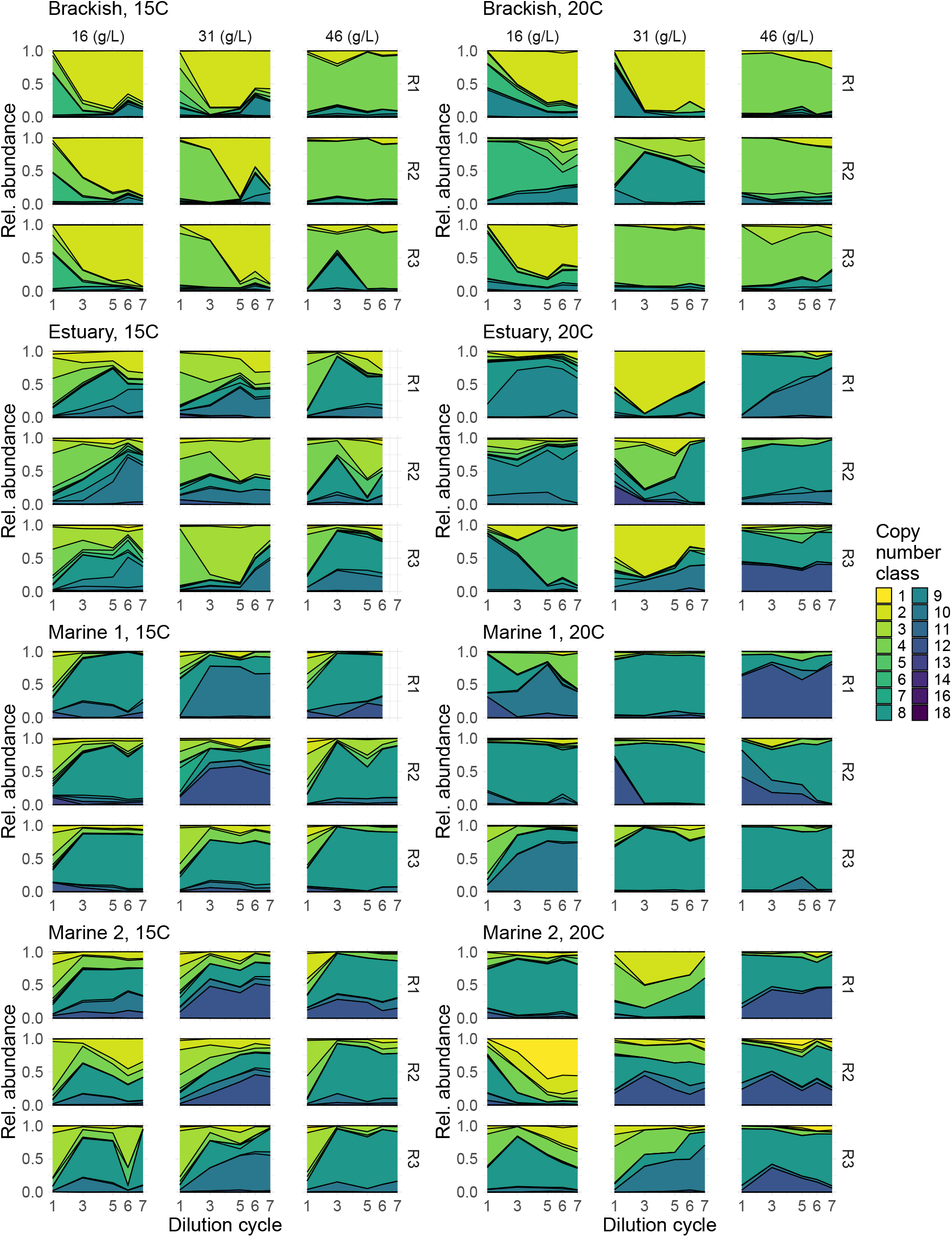
Community composition (coloured by copy number class) over 7 growth-dilution cycles. ASVs are mapped to their rRNA gene copy number class. Each copy number class describes the greatest integer less than or equal to the estimated rRNA gene copy number of an ASV. Communities are separated by the propagation temperature (columns; 15 or 20 *◦*C) and the source community (rows; Brackish, Estuary, Marine 1, Marine 2). Three replicate communities (R1-3) were propagated for each condition.

**Figure S4:**
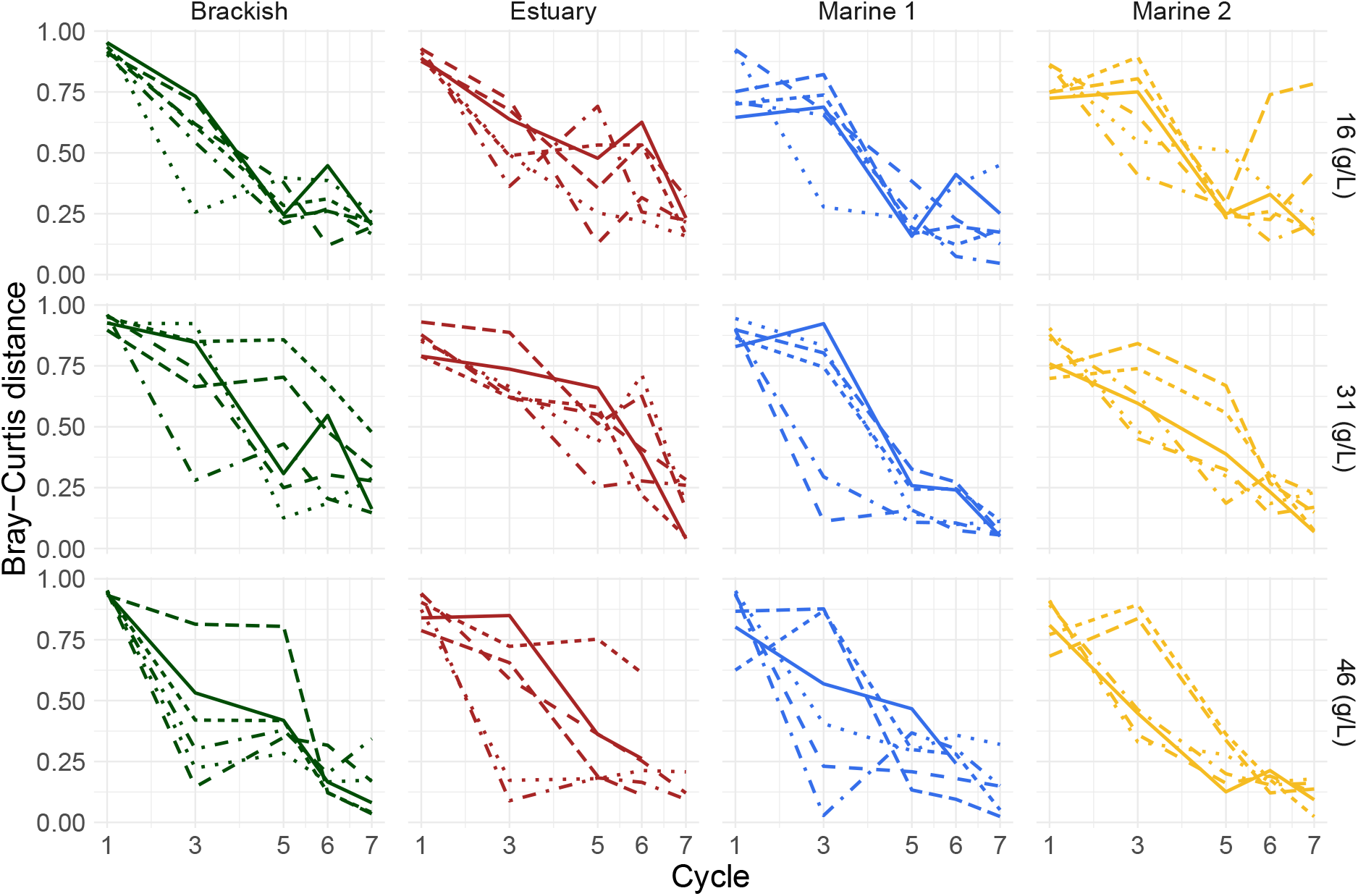
Pairwise Bray-Curtis distances between communities on subsequent days. For each starting community/salinity pair, the 3 biological replicates at two propagation temperatures are indicated with different line styles. The value plotted on cycle x represents the Bray-Curtis distance between the community on that day and the preceding cycle for which we measured community composition (e.g. the value for cycle 1 represents the distance between C0 and C1).

**Figure S5:**
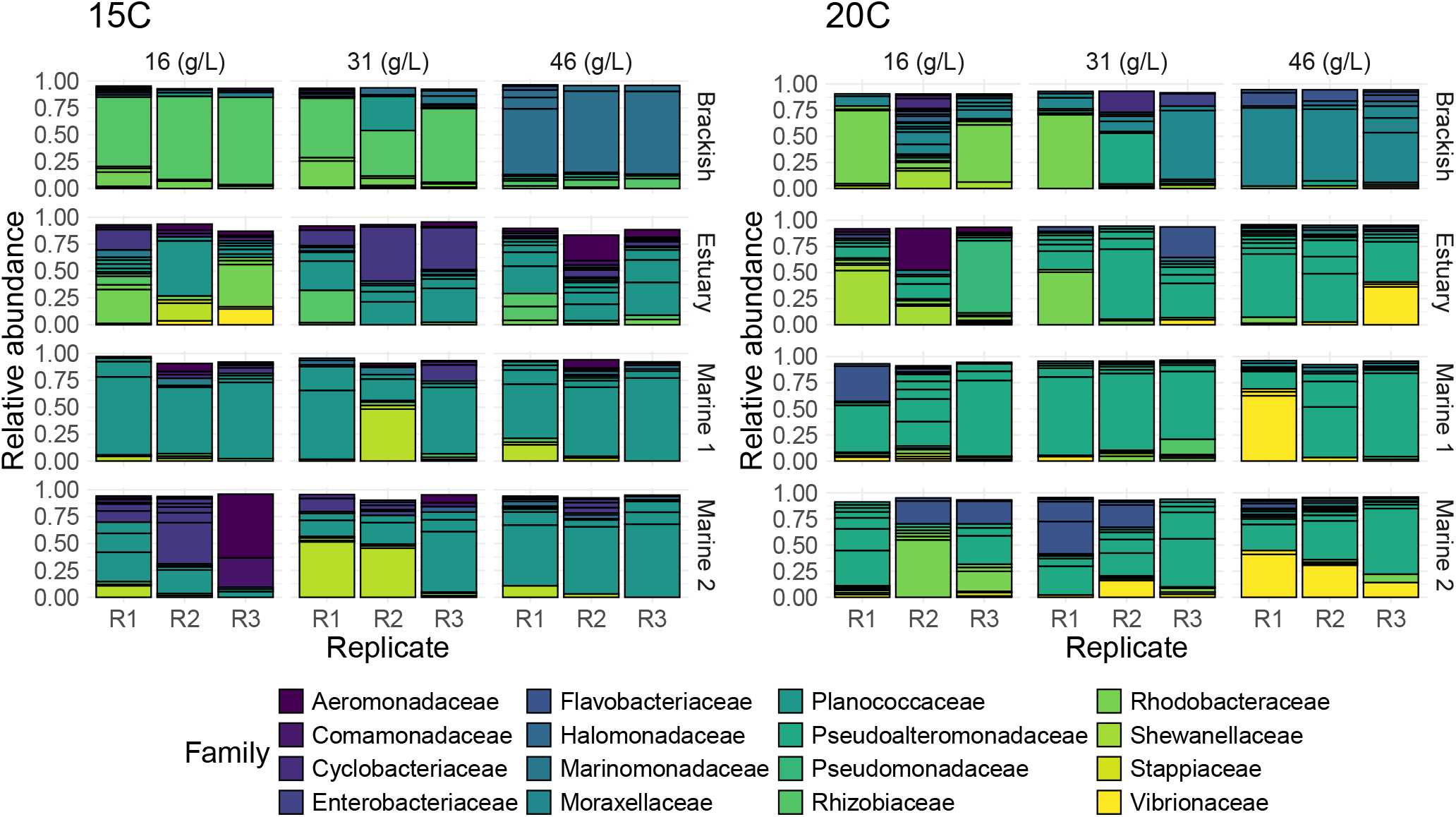
Community composition after cycle 6. Communities are separated by the temperature (left and right panel; 15*◦*C and 20*◦*C) and salinity (columns within each panel; 16, 31, or 46 g/L) they were propagated at, as well as the source community (rows within each panel; Brackish, Estuary, Marine 1/2). Three replicate communities (R1-3) were propagated for each condition. ASVs are colored by taxonomic family, and only ASVs at greater than 1% relative abundance are depicted.

**Figure S6:**
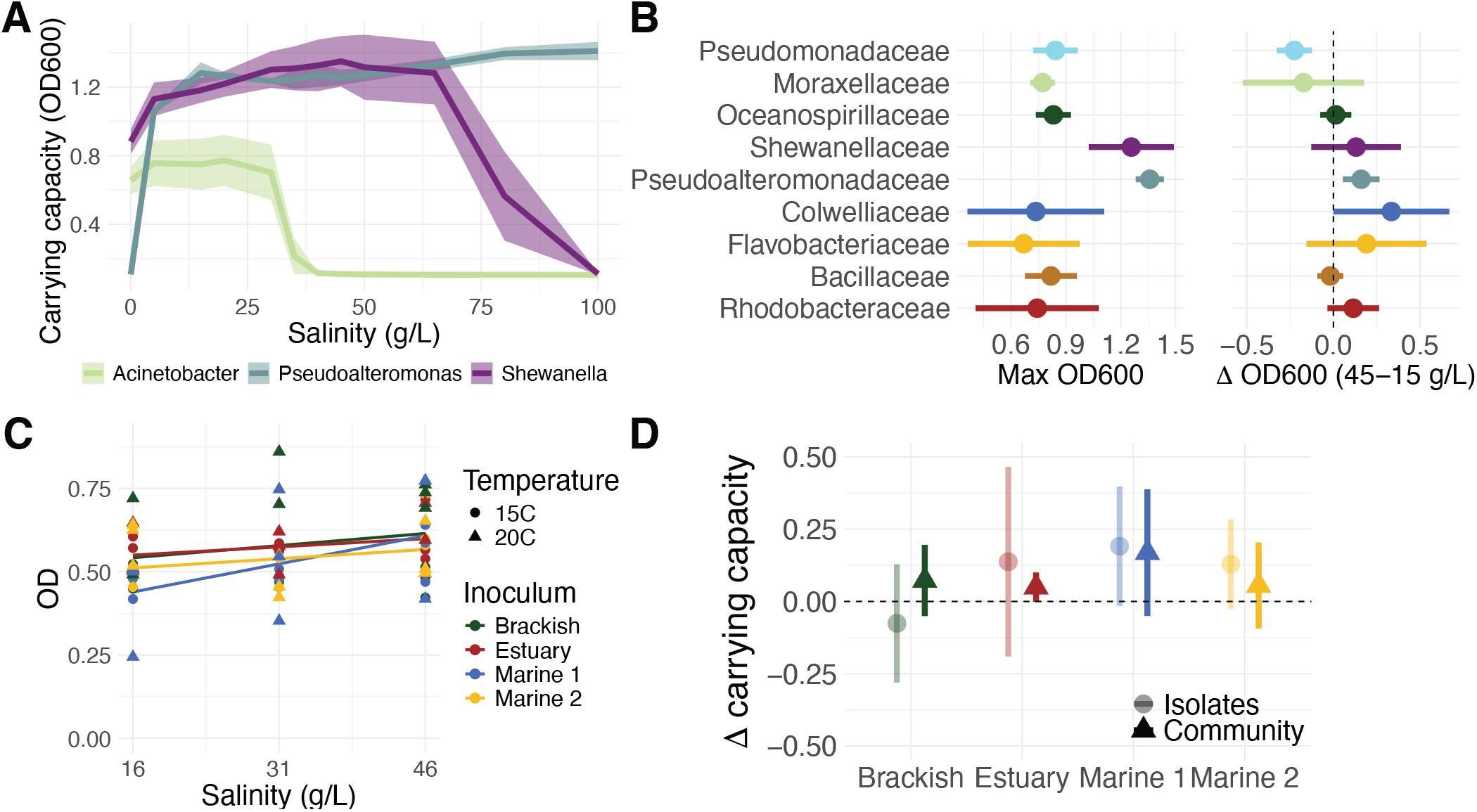
The carrying capacities of isolates and communities are robust to an increase in salinity. **A)** Measured carrying capacities as a function of salinity for the three isolates shown in Fig. 1B (Shewanella in purple, Acinetobacter in light-green, Pseudoalteromonas in blue; *n* ≥ 3 replicates). **B)** Summary statistics of isolate carrying capacities, grouped by family. Shown are the maximum carrying capacity (approximated by max OD600 in monoculture) obtained across all salinities, as well as the change in carrying capacity between growth at 45 and 15 g/L. **C)** The OD600 of each community at the end of the serial dilution experiment (C6). **D)** The mean change in carrying capacity between 15 and 45 g/L, for communities (dark triangles) and isolates (light circles).

**Figure S7:**
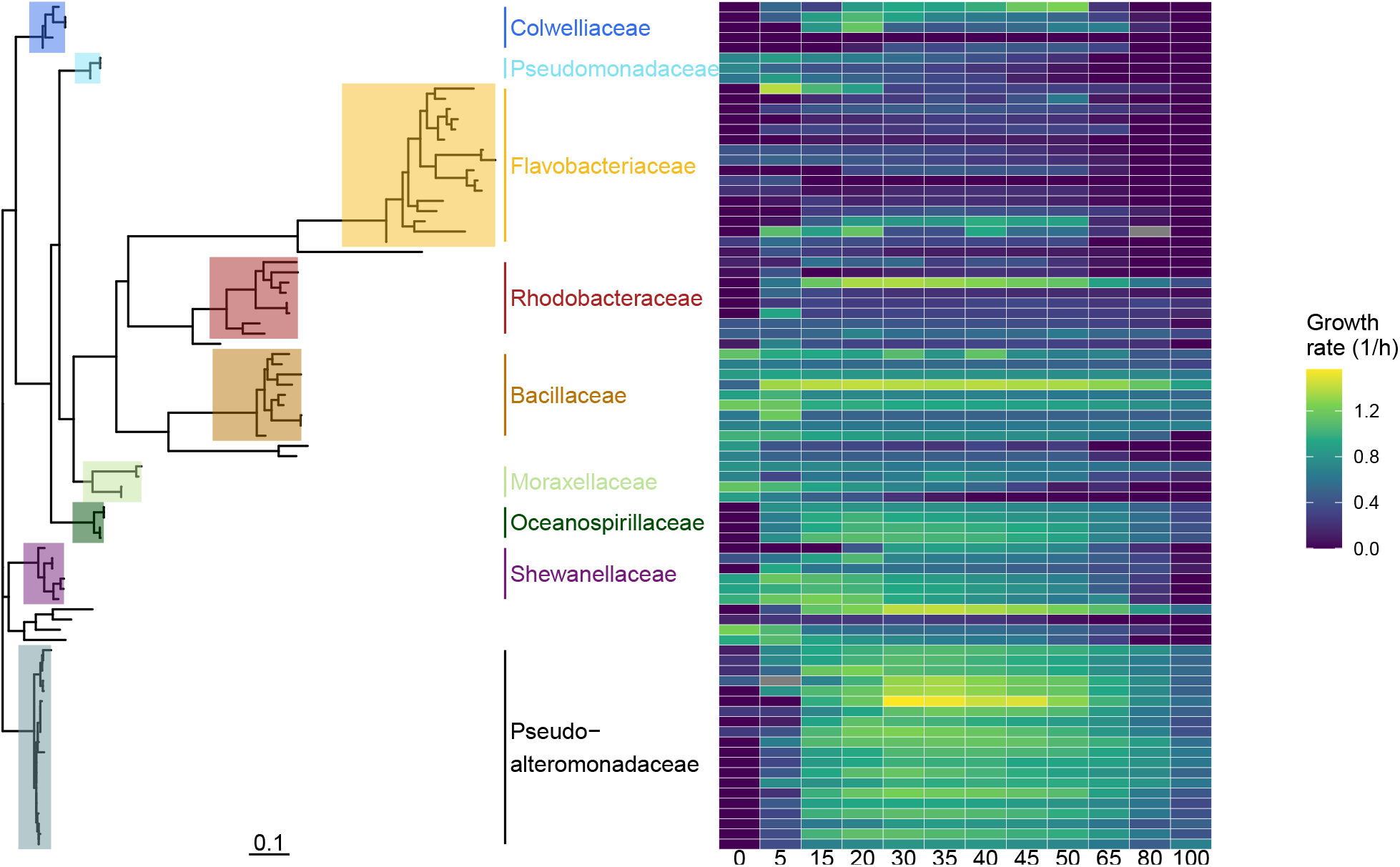
Maximum likelihood phylogenetic tree of the isolate 16S gene sequence, with the measured growth rates as a function of salinity (g/L). The tree is rooted at an arbitrary point to highlight major clades. Families with at least 3 isolates are highlighted in colors corresponding to Fig. 1C.

**Figure S8:**
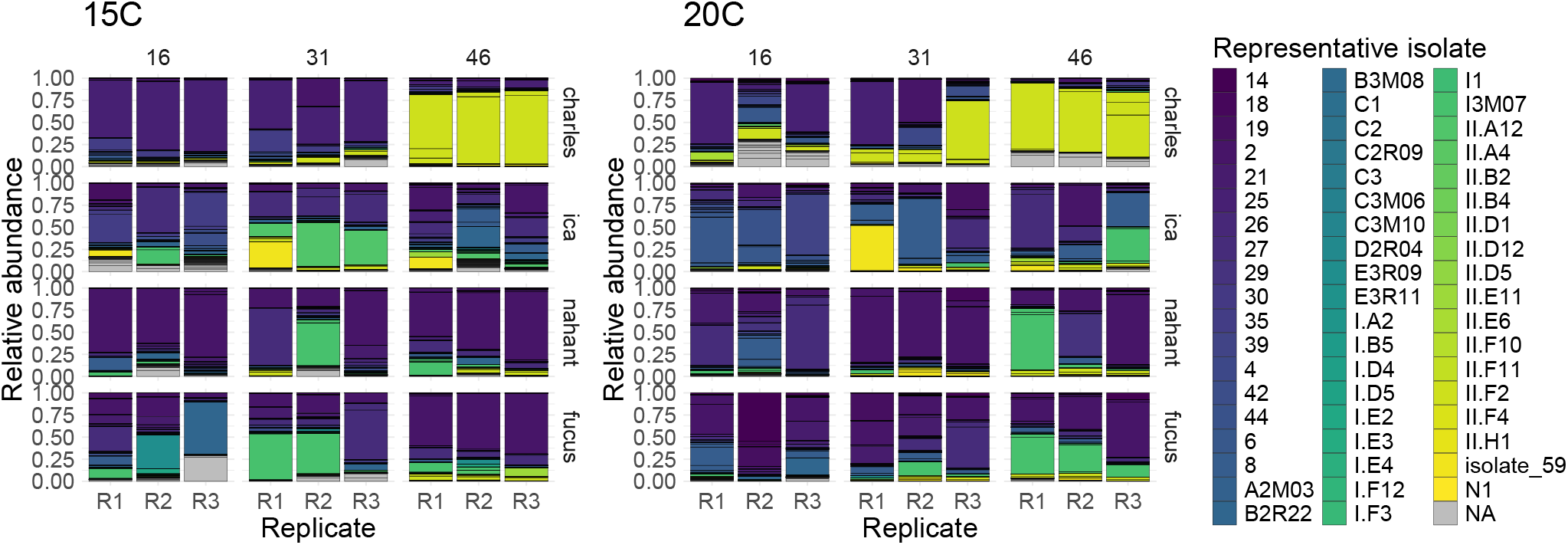
Our isolates cover the diversity observed at the end of the serial dilution experiment. An isolate is considered a match to an ASV if the 16S sequences are *>* 97% similar. If more than one isolate match the same ASV equally well, their growth rates are averaged. Columns correspond to different salinities (16, 31, or 46 g/L), subdivided into three replicates (R1, R2, R3). Rows correspond to different inoculum communities.

**Figure S9:**
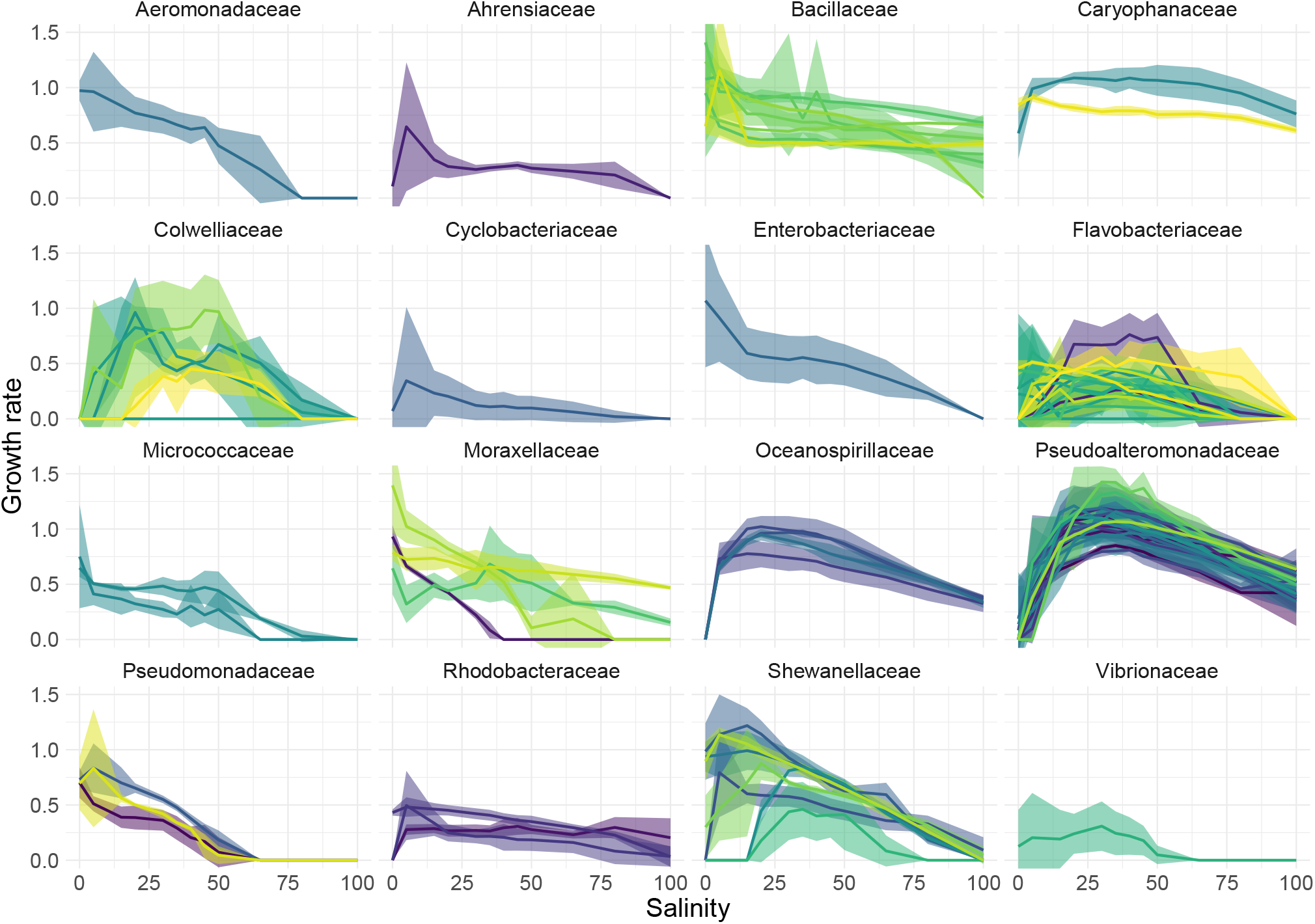
Salinity performance curves for all isolates in our dataset, grouped by family. Maximum growth rate was measured at 12 different salinities (0, 5, 15, 20, 30, 35, 40, 45, 50, 65, 80, 100 g/L). Isolates are assigned different colors, solid lines indicate the mean across all measurements for an isolate (*n* ≥ 3) and ribbons indicate mean *±* sd.

**Figure S10:**
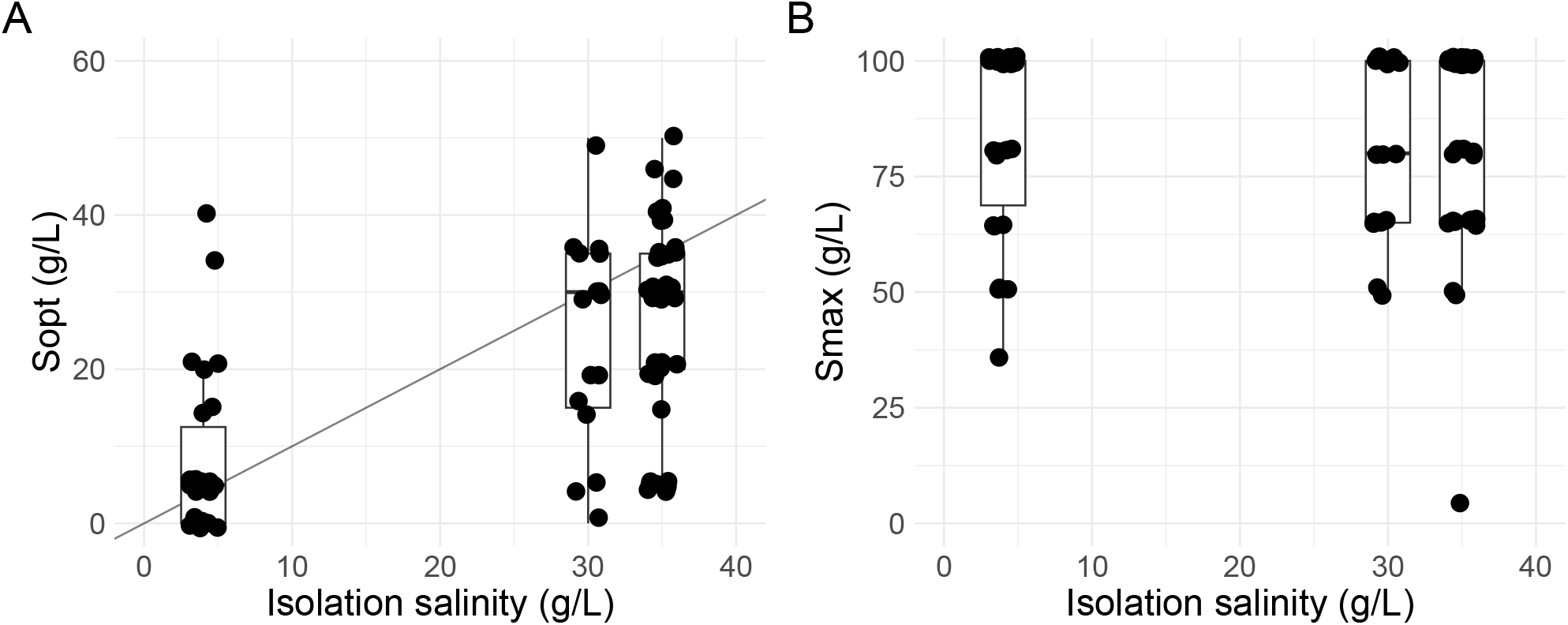
The optimal salinity for growth (Sopt) is higher for strains isolated at higher environmental salinity. A) The salinity at which an isolate reaches its maximal growth rate (Sopt) plotted as a function of the salinity of the isolation environment. B) The maximal salinity at which we observed non-zero growth for an isolate (Smax) plotted as a function of the salinity of the isolation environment. No measurements were made above 100 g/L, so many strains are at the limit of detection.

**Figure S11:**
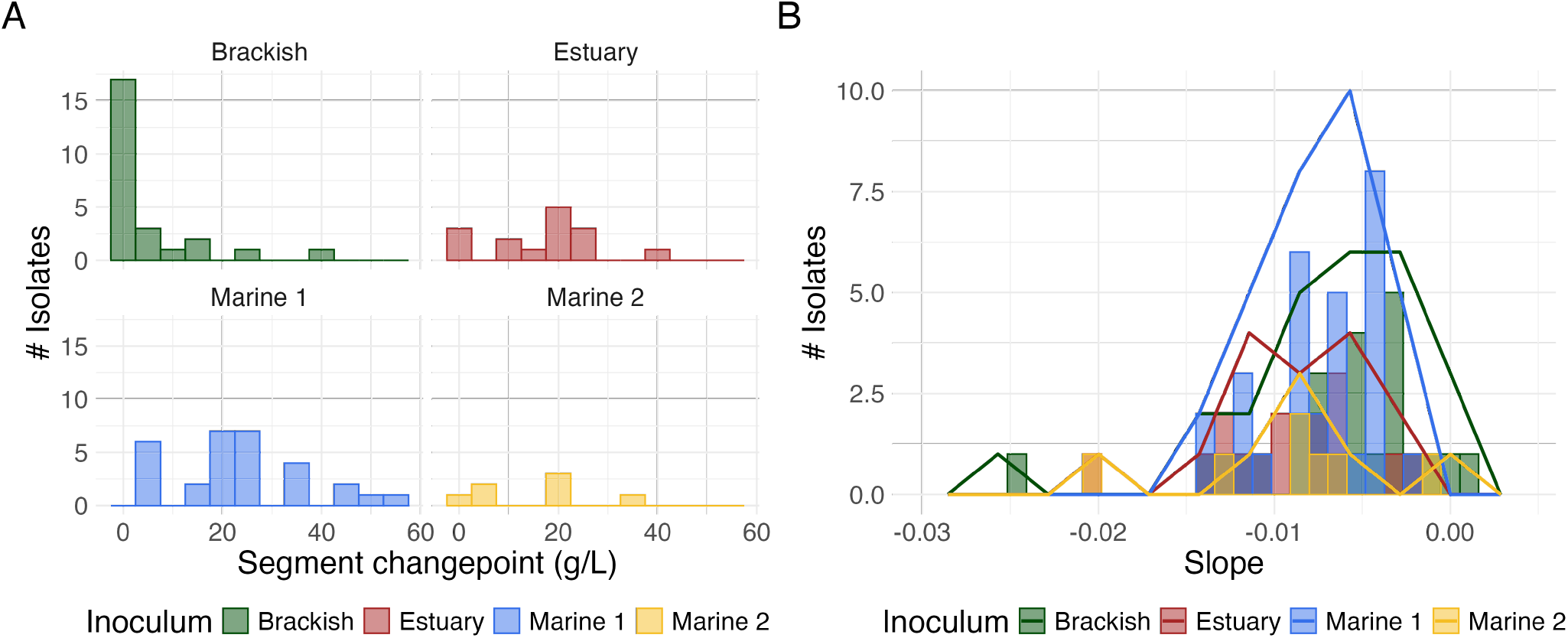
Growth parameters of marine species as fitted with 2-segment models. **A** Salinity at which the maximal growth rate is reached. This corresponds to either the breakpoint of the 2-segment model or a salinity of 0 g/L if both the first and second slope were negative. **B** The inferred slope of growth rate (1/h) as a function of salinity (g/L) after the optimum.

**Figure S12:**
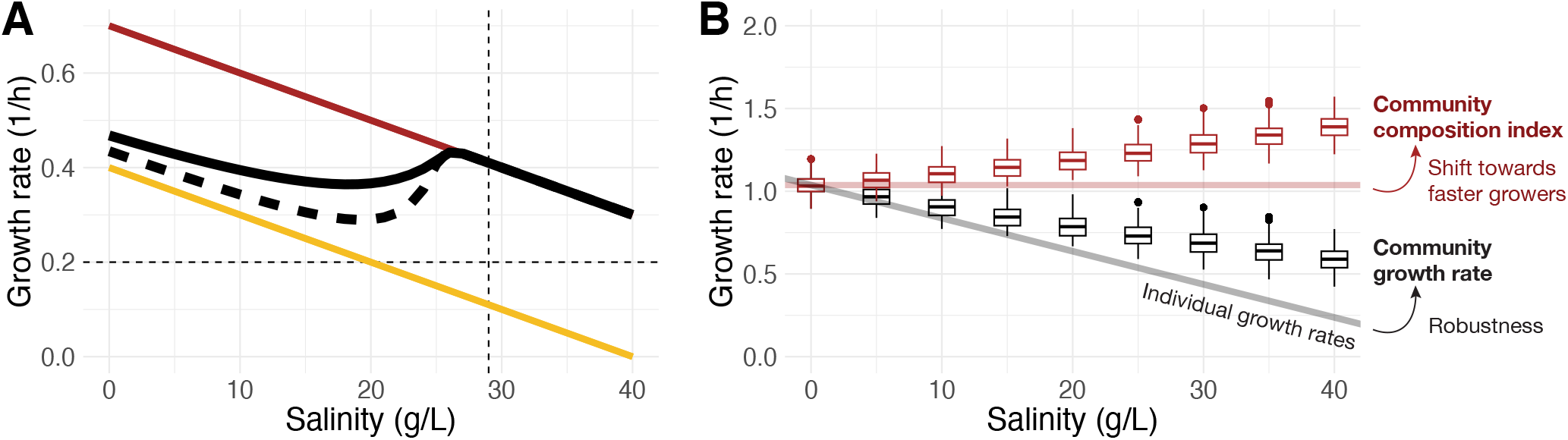
A model with daily dilution predicts that an increase in salinity shifts community composition towards faster growing species and confers robustness to the community growth rate in A) the two-species, and B) the 50-species case. A) Growth rates of the two competing species are assumed to decline at the same rate (*b* = −0.02; thin colored lines). Both a model with continuous removal (solid black line) and with daily dilution (dashed black line) show that the community growth rate declines more slowly. The dashed horizontal line indicates the strength of the removal rate (*δ* = 0.2). The dashed vertical line indicates the salinity at which the faster growing species first excludes the slower grower. B) Results of the model with daily dilution for a 50 species community. The community growth rate is robust (i.e. declines less fast than the individual species growth rates) across 50 simulations of the 50-species communities (black boxplots), while simultaneously the community composition shifts towards faster growing species (community composition index; red boxplots).

**Figure S13:**
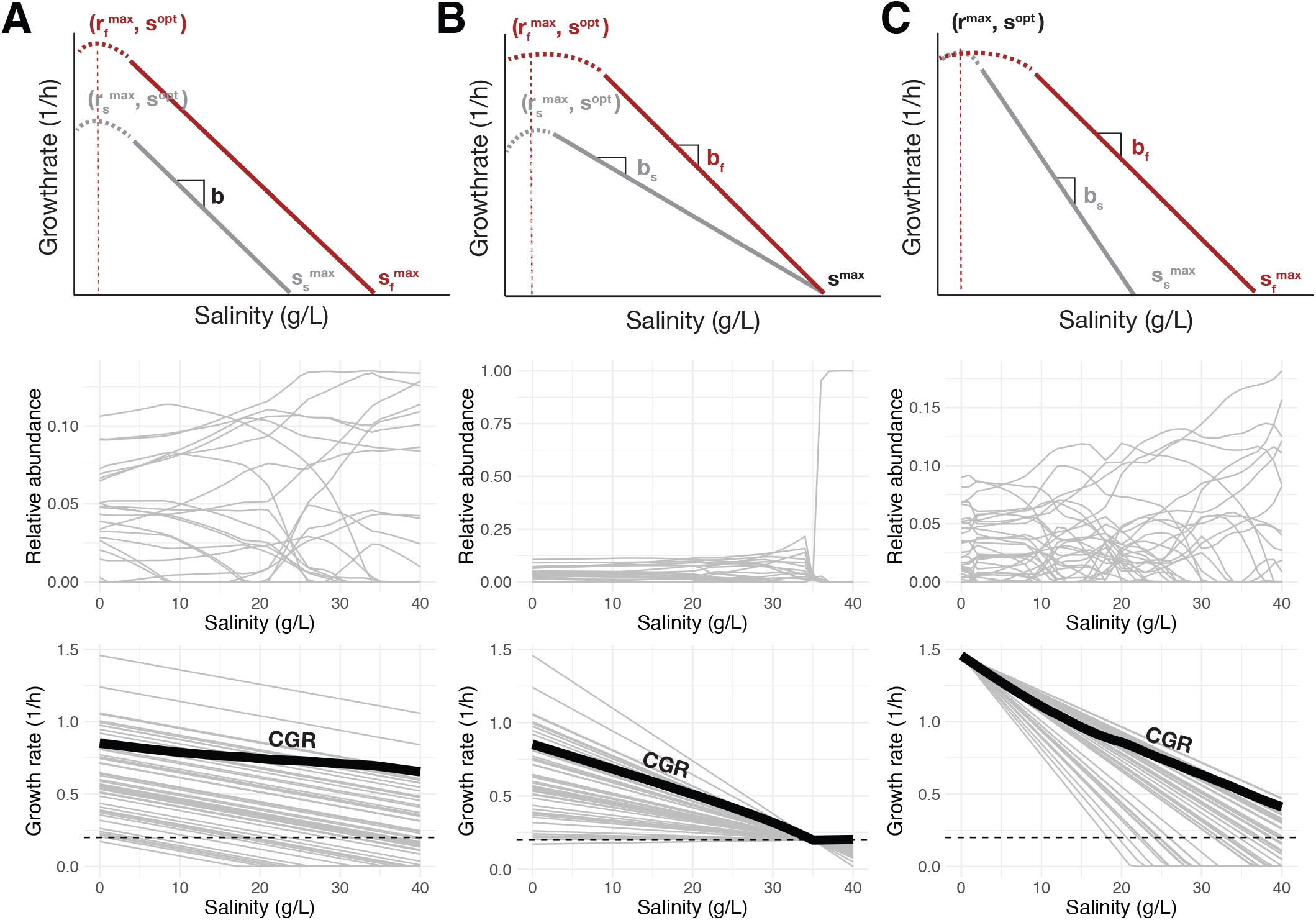
Simulations show that the proportion of faster growing species robustly increases, independent of the distribution of growth rates as a function of salinity. We compare three different scenarios: column A corresponds to a community with parallel declines in growth rate (the scenario depicted in main text Fig. 2); B corresponds to a scenario with different starting growth rate and slopes, but the same maximal salinity conducive of growth; C corresponds to a scenario with the same starting growth rate, but different slopes and different maximal salinities. In the growth rate panels (bottom row), the thick black line indicates the community growth rate (CGR), the abundance weighted mean of the realized growth rates of all species in the community. The dashed horizontal line indicates the strength of the removal rate *δ*.

**Figure S14:**
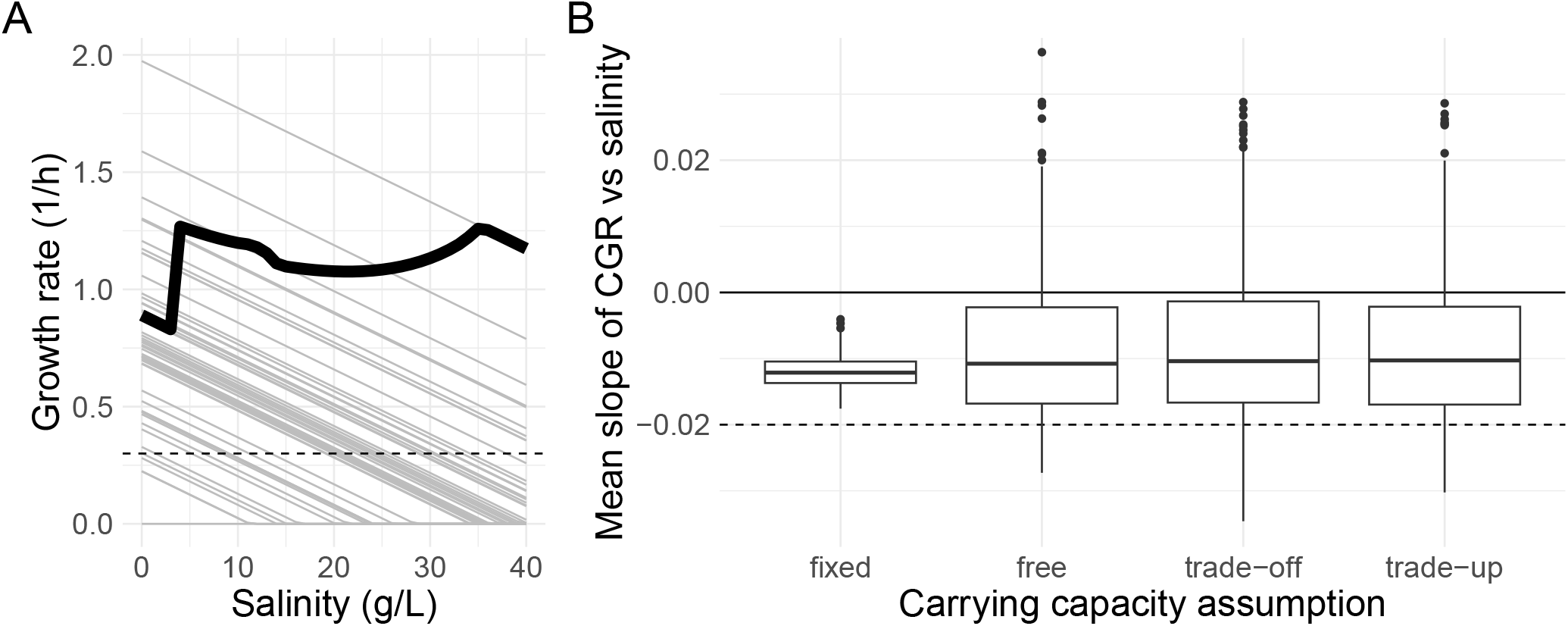
Models with alternative modeling assumptions for the carrying capacity also demonstrate robustness of the community growth rate with increasing salinity. A) Growth rates of the fifty competing species are assumed to decline at the same rate (*b* = −0.02; thin lines). If the carrying capacity varies between species, the community growth rate jumps around declines more slowly. The dashed horizontal line indicates the strength of the removal rate (*δ* = 0.2). The dashed vertical line indicates the salinity at which the faster growing species first excludes the slower grower. B) Results of 500 simulations where either all *a*_*ii*_ are fixed to 1 (‘fixed’), uniformly sampled from [0, 0.5] (‘free’), uniformly sampled from [0, 0.5] and then reordered such that the fastest growers have the highest *a*_*ii*_ (i.e. the lowest carrying capacity; ‘trade-off’), or uniformly sampled from [0, 0.5] and then reordered such that the fastest growers have the lowest *a*_*ii*_ (i.e. the highest carrying capacity; ‘trade-up’). Slopes of the CGR were estimated by fitting a linear model to the CGR at each salinity between 0 to 40 g/L. The community growth rate is robust (i.e. declines less fast than the slope of the individual species growth rates, −0.02) across all four scenarios.

**Figure S15:**
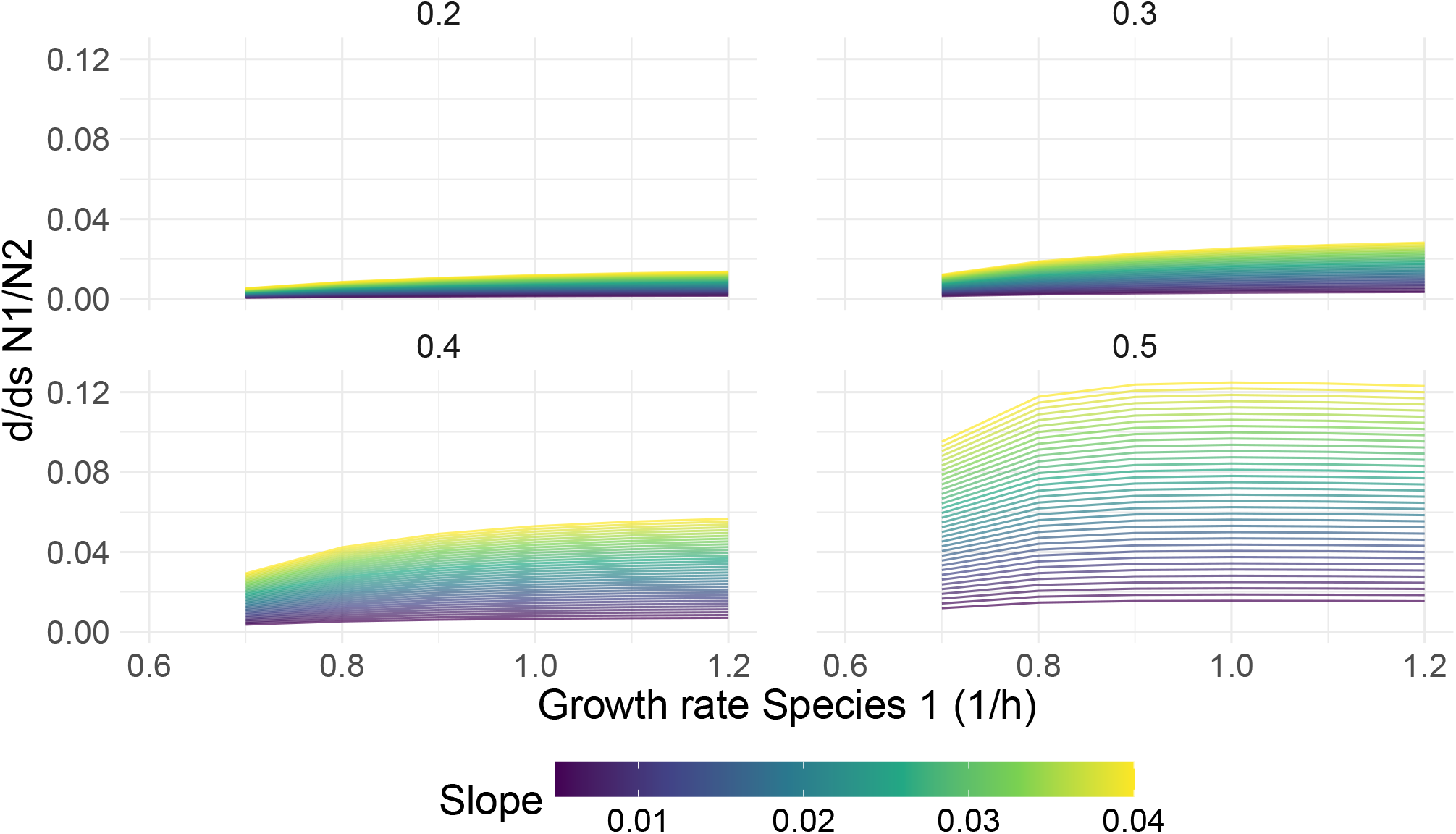
Analytical results of equation 24 for different growth rates, mortality rates (facets) and slopes of growth as a function of salinity (colors). Derivatives were calculated at the optimal salinity (here *S*_*opt*_ = 0), assuming species 2 has a growth rate *r*_*s*_ = 0.6.

**Figure S16:**
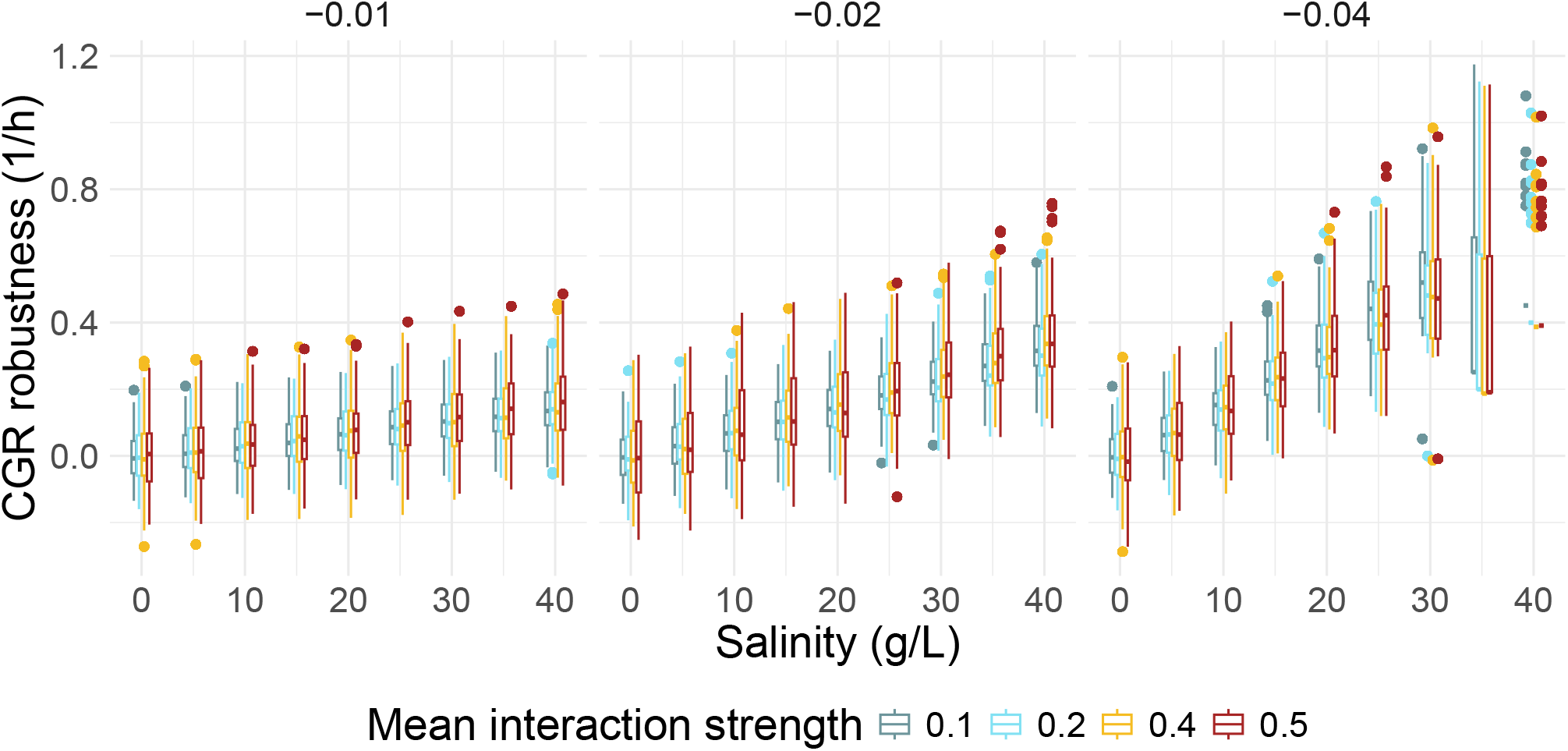
Slope of the growth rate, but not interaction strength, affect the robustness of the community growth rate (CGR) as a function of salinity. Robustness was calculated as the realized CGR - the average decline in species growth at that salinity. In these simulations, the robustness in the CGR exactly mirrors the shift in the community composition index (CCI). Columns indicate different slopes of the growth rate as a function of salinity, and colors different values of the mean interaction strength *α*. Each boxplot summarizes 50 simulations of 50-species communities.

**Figure S17:**
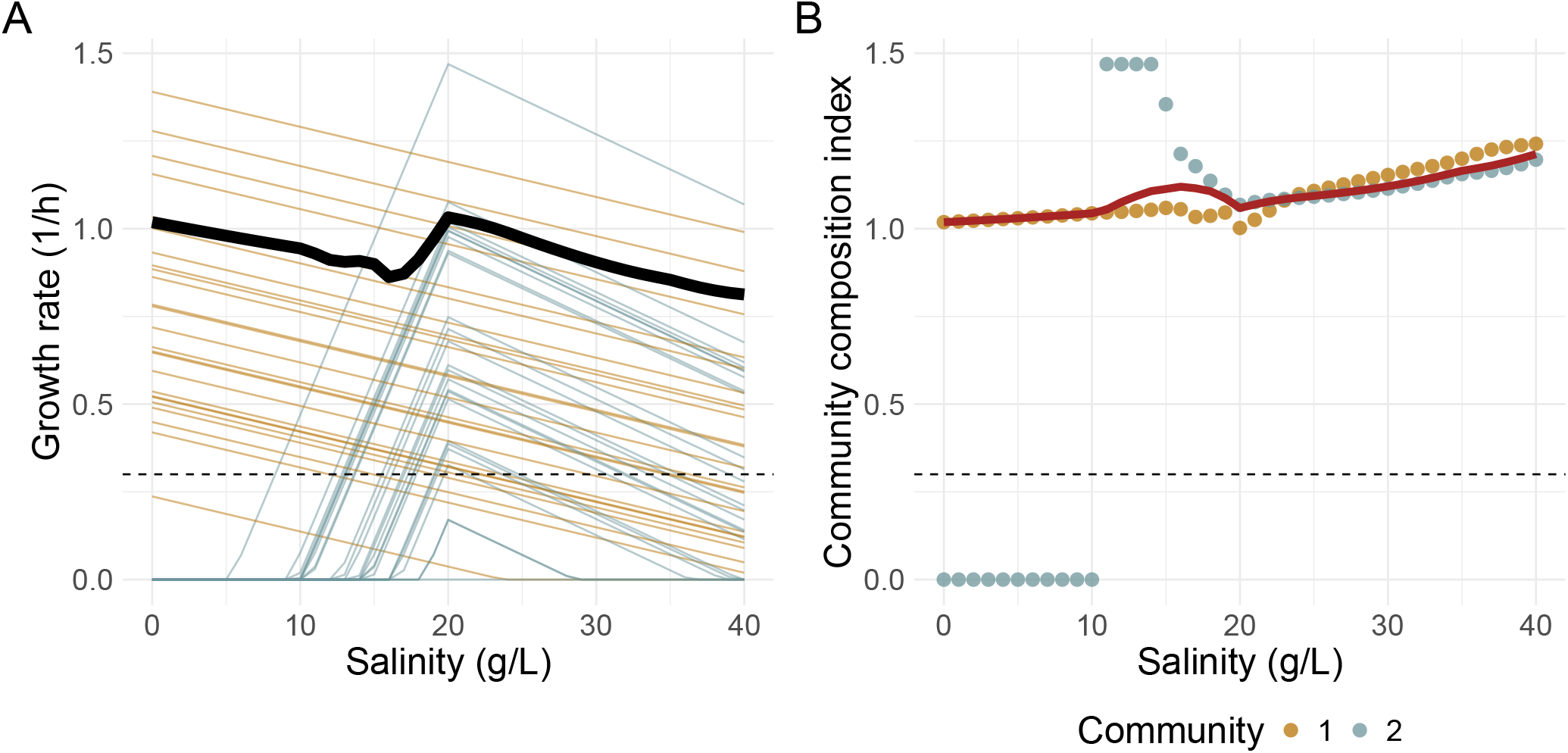
Modeling suggests that the community growth rate remains robust when isolates with two different optimal salinities are mixed. Simulations were initiated with a total of 50 isolates, half of which had *s*^*opt*^ = 0 (Community 1, yellow) and the other half *s*^*opt*^ = 20 (Community 2, blue). Community growth rate was robust across a broad range (black line), and the community composition index (red line) increased.

**Figure S18:**
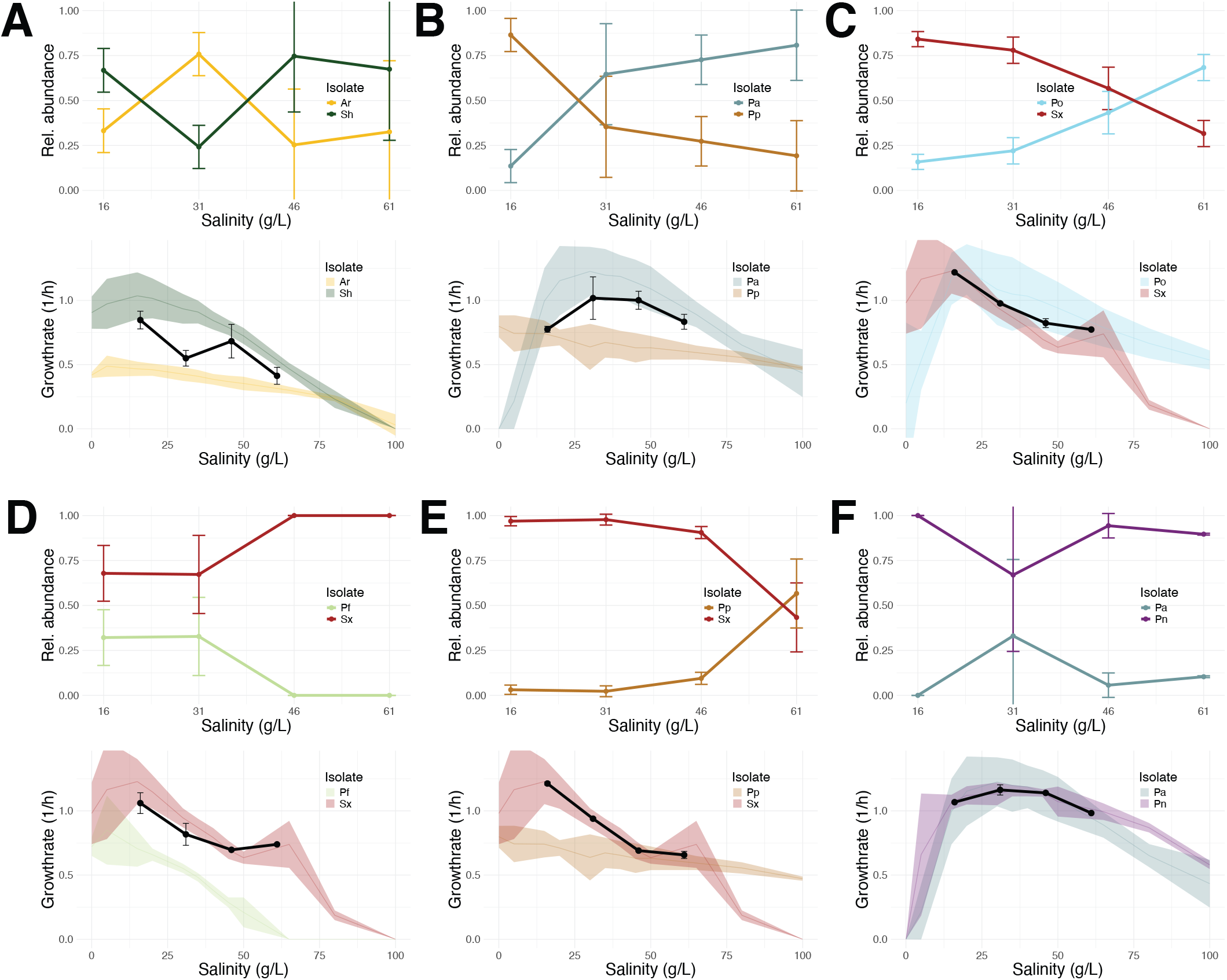
In pairwise co-culture, higher salinity favors the faster growing species. A) Salinity performance curves for the two species in each pair (ribbons show mean *±* sd across *n* ≥ 3 replicates). The black line indicates the CGR of the pairs after 14 days of propagation at 16, 31, 46, 61 g/L salinity. B) Relative abundance of both isolates in a pair after propagation at 4 salinities (16, 31, 46, 61 g/L) for 14 days. Mean and sd across three biological replicates at three different initial ratios (5:95, 50:50, 95:5). Abbreviations indicate: *Pseudoalteromonas arctica* (*Pa*), *Shewanella xiamensis* (*Sx*), *Shewanella sp*. (*Sh*), *Albirhodobacter sp*. (*Ar*), *Pseudoalteromonas ostrae* (*Po*), *Pseudoalteromonas nitrifaciens* (*Pn*), *Pseudomonas fragi* (*Pf*), and *Psychrobacter piscatorii* (*Pp*).

**Figure S19:**
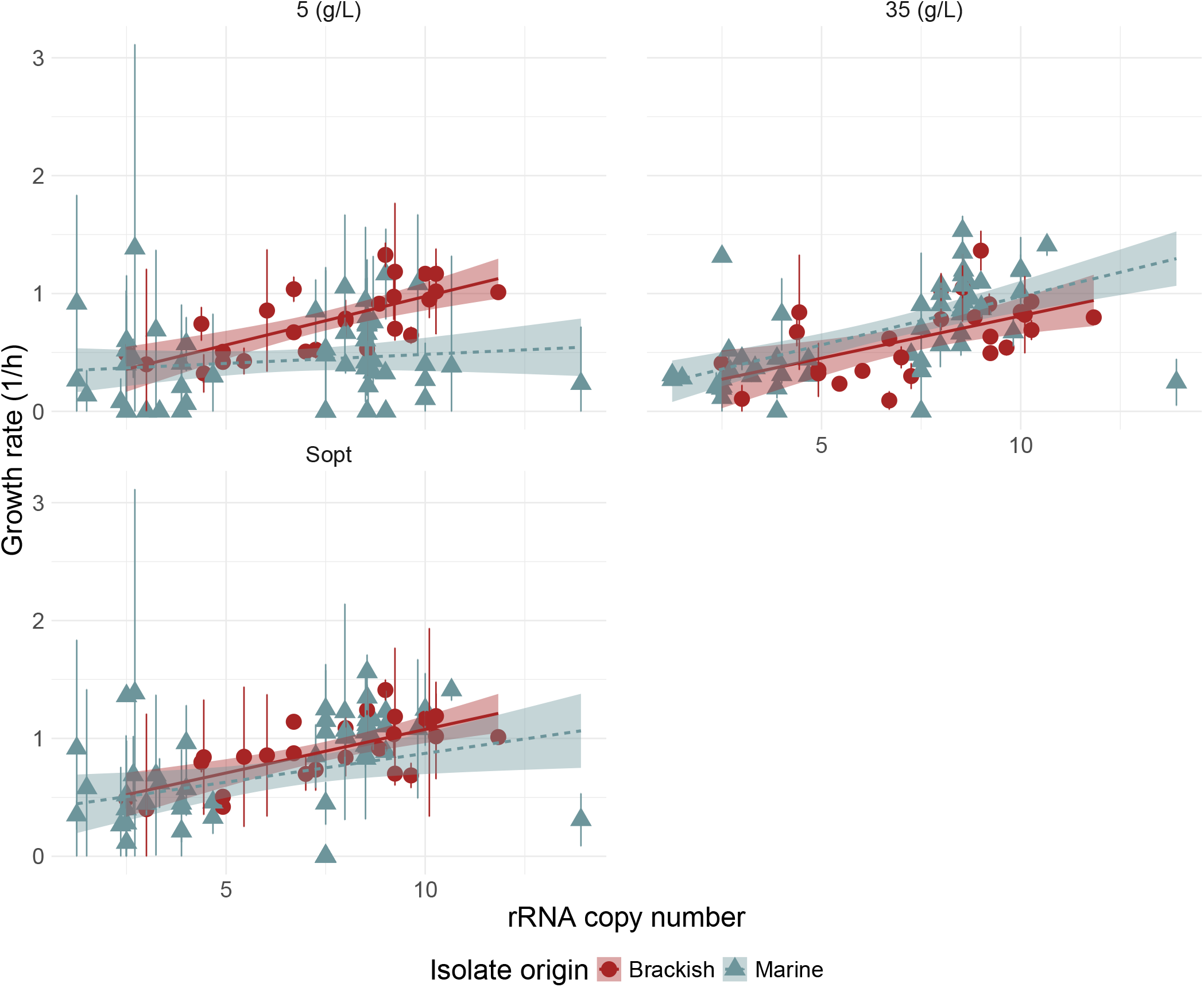
Growth rate of an isolate at its isolation salinity correlates strongly with its predicted 16S rRNA copy number. Isolates are colored by their isolation location. For the purpose of this analysis isolates from the Estuary sampling location (30 g/L) were grouped with the Marine isolates (35 g/L). The third panel, Sopt, associates each isolate with its maximal growth rate (at salinity *s*_*opt*_).

**Figure S20:**
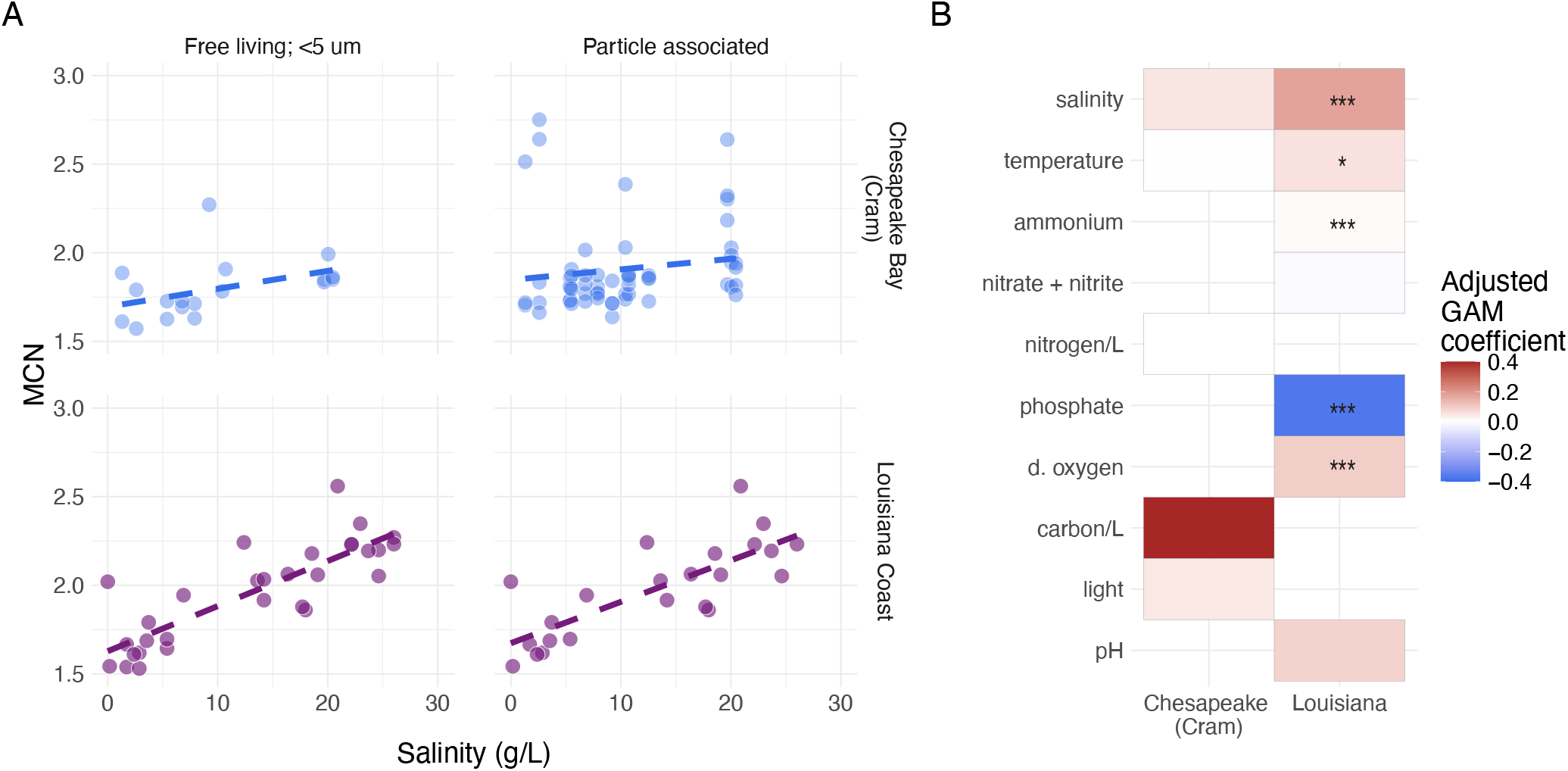
Higher salinity favors faster growers in free-living estuarine communities. A) Mean copy number of the 16S rRNA gene (MCN) of each community sampled in either the Chesapeake Bay dataset of Cram *et al*. [18] (dark blue), or the Louisiana coast dataset (purple), plotted as a function of salinity at the location and time of sampling and split by the size fraction of the communities. For the Chesapeake Bay dataset we combined both the 0.2*µm* −1.2*µm* and 1.2*µm* −5*µm* size fractions into the ‘free-living’ category, while for the Louisiana dataset this refers to the fraction 0.2*µm* −2.7*µm*. B) Effect sizes of the different predictors of mean copy number. The color corresponds to the estimated parametric coefficient in the model, multiplied by the standard deviation of the environmental predictor (values below −0.4 were assigned the same color). Environmental predictors were abbreviated with “p.” for particulate, and “d.” for dissolved. The asterisks indicate significance at \*\*\**p <* 0.001, \*\**p <* 0.01, \**p <* 0.05.

## S3 Supplementary Tables

**Table S3:**
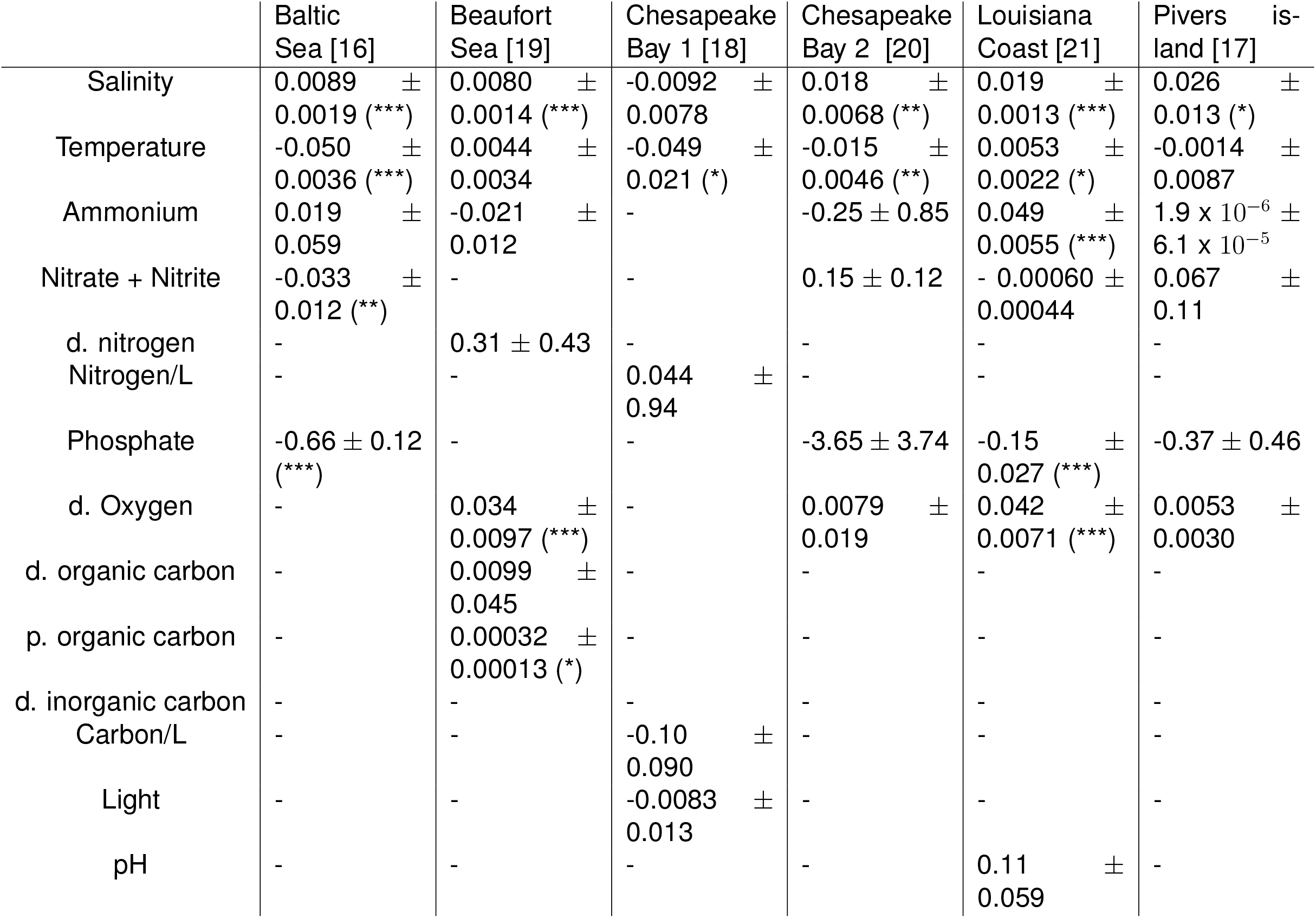
Coefficients estimated in the GAM analysis. The asterisks indicate significance at \*\*\**p <* 0.001, \*\**p <* 0.01, \**p <* 0.05. Environmental predictors were abbreviated with “p.” for particulate, and “d.” for dissolved. Values are reported *±* standard error. In the Chesapeake Bay 1 and Louisiana coast datasets, all size fractions were analyzed together.

## References

[1] Naomi M Levine, Harriet Alexander, Erin M Bertrand, Victoria J Coles, Stephanie Dutkiewicz, Suzana G Leles, and Emily J Zakem. Microbial ecology to ocean carbon cycling: from genomes to numerical models. Annual Review of Earth and Planetary Sciences, 53, 2025.

[2] Frances Spragge, Erik Bakkeren, Martin T Jahn, Elizete BN Araujo, Claire F Pearson, Xuedan Wang, Louise Pankhurst, Olivier Cunrath, and Kevin R Foster. Microbiome diversity protects against pathogens by nutrient blocking. Science, 382(6676):eadj3502, 2023.

[3] Joshua E Goldford, Nanxi Lu, Djordje Bajić, Sylvie Estrela, Mikhail Tikhonov, Alicia Sanchez-Gorostiaga, Daniel Segrè, Pankaj Mehta, and Alvaro Sanchez. Emergent simplicity in microbial community assembly. Science, 361(6401):469–474, 2018.

[4] Simon Lax, Clare I. Abreu, and Jeff Gore. Higher temperatures generically favour slowergrowing bacterial species in multispecies communities. Nature Ecology and Evolution, 4:560–567, 4 2020.

[5] Clare I. Abreu, Martina Dal Bello, Carina Bunse, Jarone Pinhassi, and Jeff Gore. Warmer temperatures favor slower-growing bacteria in natural marine communities. Science Advances, 9, 2023.

[6] Clare I. Abreu, Jonathan Friedman, Vilhelm L. Andersen Woltz, and Jeff Gore. Mortality causes universal changes in microbial community composition. Nature Communications, 10, 12 2019.

[7] Ashley Shade, Hannes Peter, Steven D Allison, Didier L Baho, Mercè Berga, Helmut Bürgmann, David H Huber, Silke Langenheder, Jay T Lennon, Jennifer BH Martiny, et al. Fundamentals of microbial community resistance and resilience. Frontiers in microbiology, 3:417, 2012.

[8] Matt Lloyd Jones, Damian William Rivett, Alberto Pascual-García, and Thomas Bell. Relationships between community composition, productivity and invasion resistance in semi-natural bacterial microcosms. elife, 10:e71811, 2021.

[9] David Gopaulchan, Christopher Moore, Naailah Ali, Darin Sukha, Sergio Leonardo Florez González, Fabio Esteban Herrera Rocha, Ni Yang, Mui Lim, Tristan P Dew, Andrés Fernando González Barrios, et al. A defined microbial community reproduces attributes of fine flavour chocolate fermentation. Nature Microbiology, 10(9):2130–2152, 2025.

[10] Bridget B McGivern, Jared B Ellenbogen, David W Hoyt, John A Bouranis, Brooke P Stemple, Rebecca A Daly, Samantha H Bosman, Matthew B Sullivan, Ann E Hagerman, Jeffrey P Chanton, et al. Polyphenol rewiring of the microbiome reduces methane emissions. The ISME Journal, 19(1):wraf108, 2025.

[11] Itzhak Mizrahi, R John Wallace, and Sarah Moraïs. The rumen microbiome: balancing food security and environmental impacts. Nature Reviews Microbiology, 19(9):553–566, 2021.

[12] Catherine A Lozupone and Rob Knight. Global patterns in bacterial diversity, 2007.

[13] A. Smajgl, T. Q. Toan, D. K. Nhan, J. Ward, N. H. Trung, L. Q. Tri, V. P.D. Tri, and P. T. Vu. Responding to rising sea levels in the mekong delta. Nature Climate Change, 5:167–174, 1 2015.

[14] Lijing Cheng, Kevin E. Trenberth, Nicolas Gruber, John P. Abraham, John T. Fasullo, Guancheng Li, Michael E. Mann, Xuanming Zhao, and Jiang Zhu. Improved estimates of changes in upper ocean salinity and the hydrological cycle. Journal of Climate, 33:10357–10381, 12 2020.

[15] Jun Zhao, Seemanti Chakrabarti, Randolph Chambers, Pamela Weisenhorn, Rafael Travieso, Sandro Stumpf, Emily Standen, Henry Briceno, Tiffany Troxler, Evelyn Gaiser, John Kominoski, Braham Dhillon, and Willm Martens-Habbena. Year-around survey and manipulation experiments reveal differential sensitivities of soil prokaryotic and fungal communities to saltwater intrusion in florida everglades wetlands. Science of The Total Environment, 858:159865, 2 2023.

[16] Meike AC Latz, Agneta Andersson, Sonia Brugel, Mikael Hedblom, Krzysztof T Jurdzinski, Bengt Karlson, Markus Lindh, Jenny Lycken, Anders Torstensson, and Anders F Andersson. A comprehensive dataset on spatiotemporal variation of microbial plankton communities in the baltic sea. Scientific Data, 11(1):18, 2024.

[17] Christopher S Ward, Cheuk-Man Yung, Katherine M Davis, Sara K Blinebry, Tiffany C Williams, Zackary I Johnson, and Dana E Hunt. Annual community patterns are driven by seasonal switching between closely related marine bacteria. The ISME journal, 11(6):1412–1422, 2017.

[18] Jacob A Cram, Ashley Hollins, Alexandra J McCarty, Grace Martinez, Minming Cui, Maya L Gomes, and Clara A Fuchsman. Microbial diversity and abundance vary along salinity, oxygen, and particle size gradients in the chesapeake bay. Environmental Microbiology, 26(1):e16557, 2024.

[19] Colleen TE Kellogg, James W McClelland, Kenneth H Dunton, and Byron C Crump. Strong seasonality in arctic estuarine microbial food webs. Frontiers in Microbiology, 10:2628, 2019.

[20] Hualong Wang, Chuanlun Zhang, Feng Chen, and Jinjun Kan. Spatial and temporal variations of bacterioplankton in the chesapeake bay: a re-examination with high-throughput sequencing analysis. Limnology and Oceanography, 65(12):3032–3045, 2020.

[21] Michael W Henson and J Cameron Thrash. Microbial ecology of northern gulf of mexico estuarine waters. Msystems, 9(8):e01318–23, 2024.

[22] Erhard Bremer and Reinhard Krämer. Responses of microorganisms to osmotic stress. Annu. Rev. Microbiol, 73:313–334, 2019.

[23] Krzysztof T Jurdzinski, Maliheh Mehrshad, Luis Fernando Delgado, Ziling Deng, Stefan Bertilsson, and Anders F Andersson. Large-scale phylogenomics of aquatic bacteria reveal molecular mechanisms for adaptation to salinity. Science advances, 9(21):eadg2059, 2023.

[24] T. A. McMeekin, R. E. Chandler, P. E. Doe, C. D. Garland, June Olley, S. Putro, and D. A. Ratkowsky. Model for combined effect of temperature and salt concentration/water activity on the growth rate of staphylococcus xylosus. Journal of Applied Bacteriology, 62:543–550, 6 1987.

[25] David W Miles, Thomas Ross, June Olley, and Thomas A McMeekin. Development and evaluation of a predictive model for the effect of temperature and water activity on the growth rate of vibrio parahaemolyticus. International Journal of Food Microbiology, 38:133–142, 9 1997.

[26] Tim N Enke, Manoshi S Datta, Julia Schwartzman, Nathan Cermak, Désirée Schmitz, Julien Barrere, Alberto Pascual-Garía, and Otto X Cordero. Modular assembly of polysaccharide-degrading marine microbial communities. Current Biology, 29(9):1528–1535, 2019.

[27] Matti Gralka, Shaul Pollak, and Otto X Cordero. Genome content predicts the carbon catabolic preferences of heterotrophic bacteria. Nature Microbiology, 8(10):1799–1808, 2023.

[28] Joel A Klappenbach, John M Dunbar, and Thomas M Schmidt. rrna operon copy number reflects ecological strategies of bacteria. Applied and environmental microbiology, 66(4):1328–1333, 2000.

[29] Benjamin RK Roller, Steven F Stoddard, and Thomas M Schmidt. Exploiting rrna operon copy number to investigate bacterial reproductive strategies. Nature microbiology, 1(11):1–7, 2016.

[30] Steven F Stoddard, Byron J Smith, Robert Hein, Benjamin RK Roller, and Thomas M Schmidt. rrn db: improved tools for interpreting rrna gene abundance in bacteria and archaea and a new foundation for future development. Nucleic acids research, 43(D1):D593–D598, 2015.

[31] Diana R Nemergut, Joseph E Knelman, Scott Ferrenberg, Teresa Bilinski, Brett Melbourne, Lin Jiang, Cyrille Violle, John L Darcy, Tiffany Prest, Steven K Schmidt, et al. Decreases in average bacterial community rrna operon copy number during succession. The ISME journal, 10(5):1147–1156, 2016.

[32] Linwei Wu, Yunfeng Yang, Si Chen, Zhou Jason Shi, Mengxin Zhao, Zhenwei Zhu, Sihang Yang, Yuanyuan Qu, Qiao Ma, Zhili He, et al. Microbial functional trait of rrna operon copy numbers increases with organic levels in anaerobic digesters. The ISME journal, 11(12):2874–2878, 2017.

[33] Thomas P Smith, Tom Clegg, Emma Ransome, Thomas Martin-Lilley, James Rosindell, Guy Woodward, Samraat Pawar, and Thomas Bell. High-throughput characterization of bacterial responses to complex mixtures of chemical pollutants. Nature Microbiology, 9(4):938–948, 2024.

[34] Daniel R Amor and Jeff Gore. Fast growth can counteract antibiotic susceptibility in shaping microbial community resilience to antibiotics. Proceedings of the National Academy of Sciences, 119(15):e2116954119, 2022.

[35] Antonio Ventosa, Joaquín J Nieto, and Aharon Oren. Biology of moderately halophilic aerobic bacteria. Microbiology and molecular biology reviews, 62(2):504–544, 1998.

[36] Steven D Allison and Jennifer BH Martiny. Resistance, resilience, and redundancy in microbial communities. Proceedings of the National Academy of Sciences, 105(supplement 1):11512–11519, 2008.

[37] Benjamin J. Callahan, Paul J. McMurdie, Michael J. Rosen, Andrew W. Han, Amy Jo A. Johnson, and Susan P. Holmes. Dada2: High-resolution sample inference from illumina amplicon data. Nature Methods, 13:581–583, 6 2016.

[38] Christian Quast, Elmar Pruesse, Pelin Yilmaz, Jan Gerken, Timmy Schweer, Pablo Yarza, Jörg Peplies, and Frank Oliver Glöckner. The silva ribosomal rna gene database project: Improved data processing and web-based tools. Nucleic Acids Research, 41, 1 2013.

[39] Kazutaka Katoh and Daron M Standley. Mafft multiple sequence alignment software version 7: improvements in performance and usability. Molecular biology and evolution, 30(4):772–780, 2013.

[40] Alexandros Stamatakis. Raxml version 8: a tool for phylogenetic analysis and post-analysis of large phylogenies. Bioinformatics, 30(9):1312–1313, 2014.

[41] Stephen F. Altschul, Warren Gish, Webb Miller, Eugene W. Myers, and David J. Lipman. Basic local alignment search tool. Journal of Molecular Biology, 215:403–410, 1990.

[42] Jari Oksanen, Gavin L. Simpson, F. Guillaume Blanchet, Roeland Kindt, Pierre Legendre, Peter R. Minchin, R.B. O’Hara, Peter Solymos, M. Henry H. Stevens, Eduard Szoecs, Helene Wagner, Matt Barbour, Michael Bedward, Ben Bolker, Daniel Borcard, Tuomas Borman, Gustavo Carvalho, Michael Chirico, Miquel De Caceres, Sebastien Durand, Heloisa Beatriz Antoniazi Evangelista, Rich FitzJohn, Michael Friendly, Brendan Furneaux, Geoffrey Hannigan, Mark O. Hill, Leo Lahti, Cameron Martino, Dan McGlinn, Marie-Helene Ouellette, Eduardo Ribeiro Cunha, Tyler Smith, Adrian Stier, Cajo J.F. Ter Braak, and James Weedon. vegan: Community Ecology Package, 2025. R package version 2. 7-1.

[43] Michael Blazanin. gcplyr: an r package for microbial growth curve data analysis. BMC bioinformatics, 25(1):232, 2024.

[44] R Core Team. R: A Language and Environment for Statistical Computing. R Foundation for Statistical Computing, Vienna, Austria, 2023.

[45] Robert May and Angela R McLean. Theoretical ecology: principles and applications. OUP Oxford, 2007.

